# The annotation and function of the Parkinson’s and Gaucher disease-linked gene *GBA1* has been concealed by its protein-coding pseudogene *GBAP1*

**DOI:** 10.1101/2022.10.21.513169

**Authors:** Emil K. Gustavsson, Siddharth Sethi, Yujing Gao, Jonathan W. Brenton, Sonia García-Ruiz, David Zhang, Raquel Garza, Regina H. Reynolds, James R. Evans, Zhongbo Chen, Melissa Grant-Peters, Hannah Macpherson, Kylie Montgomery, Rhys Dore, Anna I. Wernick, Charles Arber, Selina Wray, Sonia Gandhi, Julian Esselborn, Cornelis Blauwendraat, Christopher H. Douse, Anita Adami, Diahann A.M. Atacho, Antonina Kouli, Annelies Quaegebeur, Roger A. Barker, Elisabet Englund, Frances Platt, Johan Jakobsson, Nicholas W. Wood, Henry Houlden, Harpreet Saini, Carla F. Bento, John Hardy, Mina Ryten

**Affiliations:** Genetics and Genomic Medicine, Great Ormond Street Institute of Child Health, University College London, London, UK; Aligning Science Across Parkinson’s (ASAP) Collaborative Research Network, Chevy Chase, MD, 20815; Astex Pharmaceuticals, 436 Cambridge Science Park, Cambridge, United Kingdom; NIHR Great Ormond Street Hospital Biomedical Research Centre, University College London, London, UK; Laboratory of Molecular Neurogenetics, Department of Experimental Medical Science, Wallenberg Neuroscience Center and Lund Stem Cell Center, Lund, Sweden; Department of Clinical and Movement Neurosciences, UCL Queen Square Institute of Neurology, University College London, London, UK; The Francis Crick Institute, London, UK; Department of Neurodegenerative Disease, UCL Queen Square Institute of Neurology, University College London, London, UK; Laboratory of Neurogenetics, National Institute on Aging, National Institutes of Health, Bethesda, MD, USA; Laboratory of Epigenetics and Chromatin Dynamics, Department of Experimental Medical Science, Lund Stem Cell Center, Lund University, Lund, Sweden; Wellcome-MRC Cambridge Stem Cell Institute and John Van Geest Centre for Brain Repair, Department of Clinical Neurosciences, University of Cambridge, Cambridge, UK; Department of Clinical Neurosciences, University of Cambridge, Clifford Albutt Building, Cambridge, UK; Department of Neuropathology, University of Lund, Lund, Sweden; Department of Pharmacology, University of Oxford, Oxford, UK; Department of Neuromuscular Disease, UCL Queen Square Institute of Neurology, UCL, London, UK; Reta Lila Weston Institute, UCL Queen Square Institute of Neurology, UCL, London, UK; UK Dementia Research Institute at UCL, UCL Queen Square Institute of Neurology, UCL, London, UK; NIHR University College London Hospitals Biomedical Research Centre, London, UK; Institute for Advanced Study, The Hong Kong University of Science and Technology, Hong Kong SAR, China

## Abstract

The human genome contains numerous duplicated regions, such as parent-pseudogene pairs, causing sequencing reads to align equally well to either gene. The extent to which this ambiguity complicates transcriptomic analyses is currently unknown. This is concerning as many parent genes have been linked to disease, including *GBA1,* causally linked to both Parkinson’s and Gaucher disease. We find that most of the short sequencing reads that map to *GBA1*, also map to its pseudogene, *GBAP1*. Using long-read RNA-sequencing in human brain, where all reads mapped uniquely, we demonstrate significant differences in expression compared to short-read data. We identify novel transcripts from both *GBA1* and *GBAP1*, including protein-coding transcripts that are translated *in vitro* and detected in proteomic data, but that lack GCase activity. By combining long-read with single-nuclear RNA-sequencing to analyse brain-relevant cell types we demonstrate that transcript expression varies by brain region with cell-type-selectivity. Taken together, these results suggest a non-lysosomal function for both GBA1 and GBAP1 in brain. Finally, we demonstrate that inaccuracies in annotation are widespread among parent genes, with implications for many human diseases.

## MAIN

The human genome contains regions that evade comprehensive analysis through short-read sequencing technologies and thus remain poorly studied. While these difficulties can be attributed to challenges with sequencing (e.g., high GC content), they are most commonly the result of duplicated genomic regions^1^. This leads to sequencing reads aligning to multiple genomic locations due to a high degree of sequence similarity, a phenomenon known as multimapping. Given that defective gene copies with high sequence similarity to their parent genes, termed pseudogenes, are frequently found in the human genome this is a common problem^2^.

While the impact of multimapping has been investigated in the context of pathogenic variant detection and can cause variants to be “missed” using conventional analyses^3^, the effect of multimapping on transcriptomic analyses has received less attention despite the problem being similar in nature^4^. This is surprising given the considerable number of genes affected, many of which are implicated in human disease. Short-read RNA-sequencing (RNA-seq) has been crucial to our understanding of transcript annotation, gene expression and its tissue and cell-type specific regulation. However, a major challenge in analysing these datasets is the difficulty of annotating parent-pseudogene pairs due to reads that cannot unambiguously map to either the parent gene or pseudogene, and so accurately quantifying gene expression.

Here, we focused on the disease-relevant example of *GBA1* and its expressed pseudogene *GBAP1*. *GBA1* encodes glucocerebrosidase (GCase), a lysosomal hydrolase^5^ that degrades the glycosphingolipid, glucosylceramide^6^. Mutations in *GBA1* result in decreased GCase activity causing Gaucher disease (GD)^7–11^ when biallelic, and when heterozygous are among the most important genetic risk factors for Parkinson’s disease (PD)^12–15^. Heterozygous *GBA1* mutations also contribute to a more rapid progression of motor and non-motor symptoms in PD^16–20^ and appear to be important predictors for non-motor symptom progression after deep brain stimulation surgery in patients with PD^21,22^.

To address the limitations of short sequencing reads, which seldom span multiple splice junctions^23^, we utilised long-read RNA-seq to examine human brain regions and iPSC-derived brain cells in depth. Our focus was on *GBA1* and *GBAP1*, and we discovered significant differences in gene expression compared to short-read RNA-seq. Moreover, we identified a significant number of novel transcripts from both genes, comprising novel protein-coding transcripts. We supported these findings by integrating short-read RNA-seq data, biochemistry, and proteomic data, which validated the novel protein-coding transcripts and confirmed that *GBAP1* is translated in cells and human brain. Furthermore, we utilized both long-read sequencing and annotation-agnostic short-read sequencing data and found that inaccuracies in annotation is common among parent genes. **Fig. 1** summarizes our analyses.

**Fig. 1:**
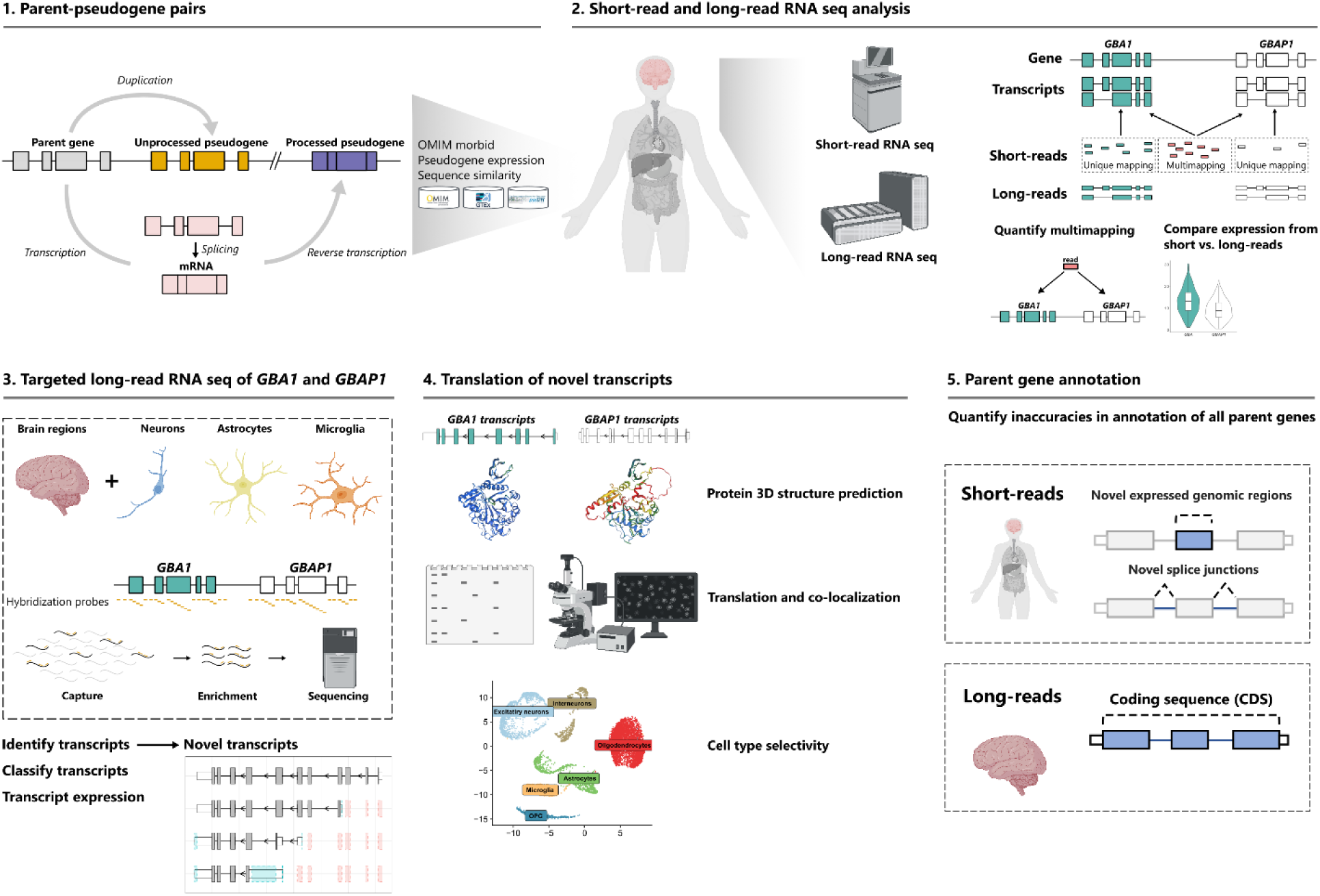
Scheme of data generation and analysis overview. Schematics outlining the methodological framework used in this study.

## RESULTS

### Pseudogenes are commonly expressed and alternatively spliced across human tissues

We started by quantifying pseudogenes from GENCODE (v38) annotation to investigate their impact on transcriptomic analyses. We identified a total of 14,709 pseudogenes in the human genome^2,24^, which can be divided into processed pseudogenes (*n* = 10,666) and unprocessed pseudogenes (*n* = 3,565), derived from retrotransposition of processed mRNAs and segmental duplications, respectively (**Fig. 2a**). To date, 10,370 pseudogenes have been confidently assigned to 3,665 unique parent genes (**Supplementary Table 1**)^25^. We found that 734 (20.0%; **Fig. 2b**) parent genes were linked to 1,015 OMIM phenotypes, accounting for 17.0% of all OMIM disease genes (https://omim.org/)^26^.

**Fig. 2:**
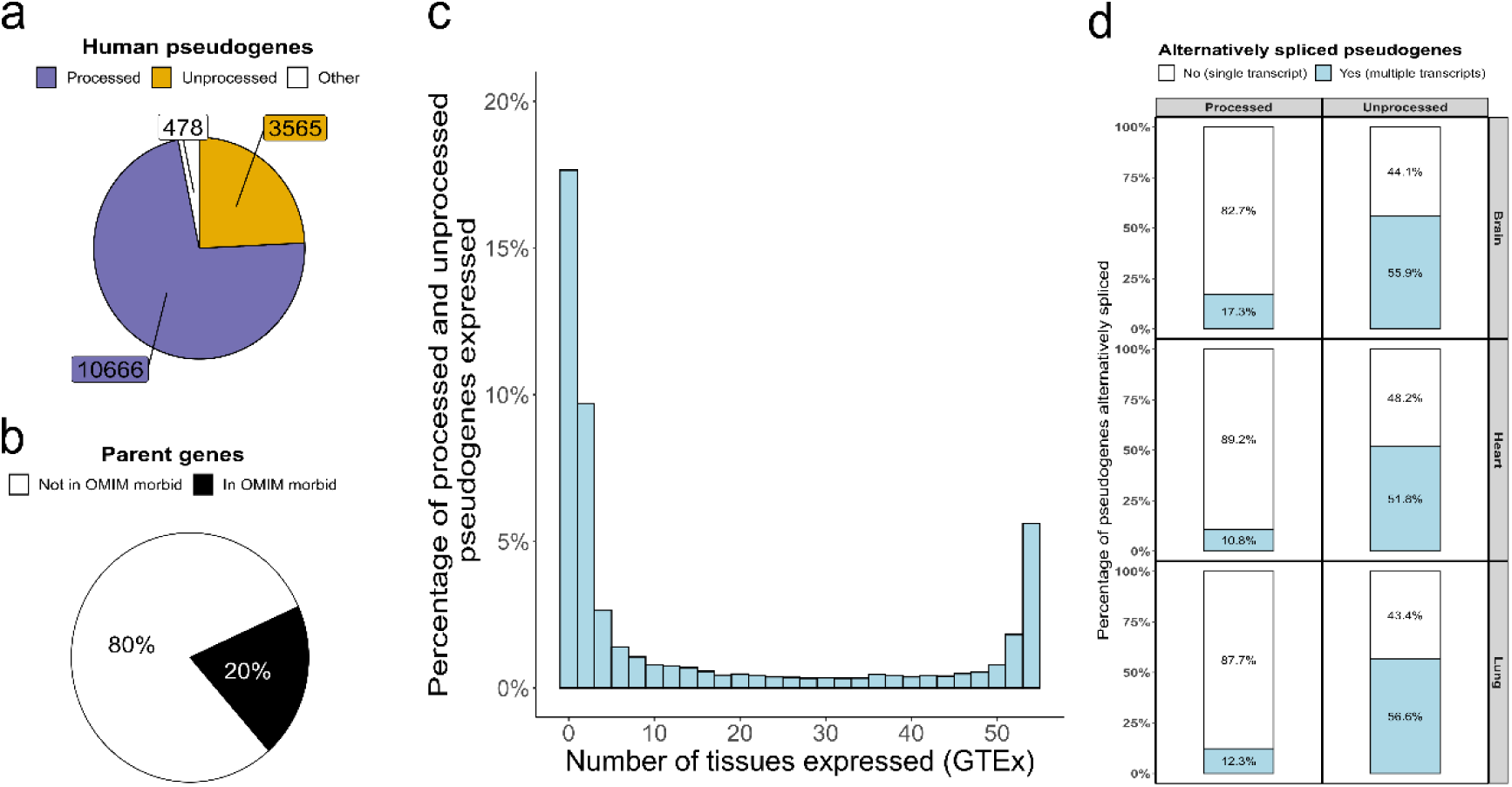
Pseudogenes are frequent and expressed across human tissues. **a,** Pie chart showing the number of annotated pseudogenes that represent processed, unprocessed, or other pseudogenes. Other pseudogenes include unitary, IG (Inactivated immunoglobulin) and TR (T-cell receptor) pseudogenes. **b,** Pie chart depicting the percentage of parent genes that are OMIM disease genes (https://omim.org). **c,** Histogram showing tissue expression of pseudogenes as assessed using uniquely mapping reads (generated by the Genotype-Tissue Expression Consortium, GTEx v8). **d,**

To examine pseudogene expression across tissues, we used uniquely mapped short-read RNA-seq data generated by the Genotype-Tissue Expression (GTEx)^27,28^ Consortium (v8, accessed 10/11/2021). We found that 64.7% of pseudogenes are expressed in ≥ 1 tissue (**Fig. 2c**), and that on average, 25.7 ± 2.5% of pseudogenes are expressed per tissue (*n* = 41; **Supplementary Fig. 1**). We then assessed the percentage of expressed pseudogenes that are alternatively spliced (> 1 transcript expressed) across human brain, heart, and lung samples using publicly available long-read RNA-seq data. On average, we found that 54.8 ± 2.6% of unprocessed pseudogenes and 13.5 ± 3.4% of processed pseudogenes are alternatively spliced (**Fig. 2d**). Taken together, this is consistent with the observation that a proportion of pseudogenes are of functional importance^29^.

### Multimapping results in significant underestimation of *GBA1* expression in human brain

We next examined the sequence similarity between pseudogenes and their parent genes as a way to investigate the potential functionality and complicating effects of the widespread expression and alternative splicing of pseudogenes. Our findings revealed that pseudogenes share an average of 80.0 ± 13.4% sequence similarity to the coding sequence (CDS) with their parent genes (**Fig. 3a**). As a result, genomic regions containing pseudogenes have the potential to confound transcriptomic analyses in all human tissues for a considerable proportion of protein-coding genes, including many that are causally linked to disease.

**Fig. 3:**
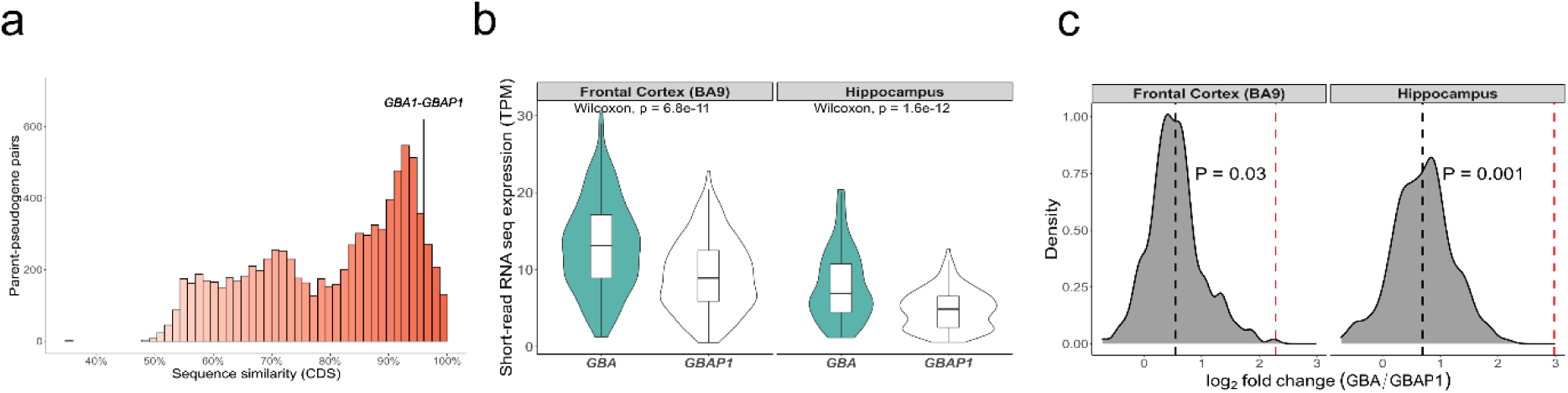
High sequence similarity causes inaccuracies in *GBA1* and *GBAP1* expression. **a,** Histogram depicting sequence similarity of parent-pseudogene pairs across coding sequences (CDS). *GBA1* and *GBAP1* 96% sequence similarity. ***b*,** Expression in transcripts per million (TPM) of *GBA1* and *GBAP1* from GTEx using gene-level expression measures (10/11/2021, v8). ***c*,** Density plot of log2 fold change of *GBA1* (numerator) and *GBAP1* (denominator) from GTEx using gene-level expression measures (10/11/2021, v8). The black dotted line represents the mean log2 fold change of *GBA1* and *GBAP1* using GTEx-derived data, while the red dotted line represents the log2 fold change generated through direct cDNA Oxford Nanopore technologies (ONT) sequencing from pooled human frontal cortex (n = 26) and hippocampus (n = 27) (total library size: 42.7 million and 48.04 million reads, respectively).

To explore this hypothesis in detail, we focused on the parent-pseudogene pair, *GBA1*-*GBAP1*^30^. This choice was driven by: (i) the high sequence similarity of *GBA1-GBAP1* of 96%, which we reasoned would make both genes prone to inaccuracies in gene expression measures and transcript annotation (**Fig. 3a**). (ii) *GBAP1*’s broad tissue expression (determined using RNA-seq data provided by GTEx), which means that simply masking its specific genomic region during mapping would be incorrect) (**Supplementary Fig. 2**). (iii) *GBA1* has been extensively studied due to its widely known role in disease, and its pseudogene is well-recognised.

We began by studying *GBA1* and *GBAP1* expression using gene-level measures from human tissues (*n* = 41) available through GTEx. Counter to previous RT-PCR-based quantifications showing that *GBA1* is expressed at significantly higher levels than *GBAP1*^31^, we found *GBA1* and *GBAP1* expression to be equivalent in many tissues (**Supplementary Fig. 3**), including the human brain (log_2_ fold change = 0.9 ± 0.5) (**Fig. 3b**). We questioned whether this observation could be explained by multimapping reads, which are often discarded in standard processing and so do not contribute to gene-level quantification of expression in many publicly available data sets (e.g., GTEx^27^, PsychENCODE^32^ and recount3^33^). To explore this question, we re-analysed publicly available short-read RNA-seq of human anterior cingulate cortex samples derived from 18 individuals, (*n,* control = 5, PD, with or without dementia = 13)^34^. Using this high-depth data set (100-bp paired-end reads, with a mean depth of 182.9 ± 14.9 million read pairs per sample), we assessed the proportion of reads that uniquely mapped to *GBA1*. We found that only 41.7 ± 11.2% of all reads mapped to *GBA1* were uniquely mapped (**Extended Data Fig. 1a**), with 96.0 ± 2.0% of multimapped reads also aligning to *GBAP1* (**Extended Data Fig. 1b**). Considering that most reads mapped to *GBA1* and *GBAP1* are not used for quantification, we concluded that long-read RNA-seq would be required to assess their relative expression. Therefore, we applied direct cDNA Oxford Nanopore sequencing (ONT) to pooled human frontal lobe (*n* individuals = 26) and hippocampus samples (*n* individuals = 27) (total library size: 42.7 million and 48.0 million reads, respectively) and found higher expression of *GBA1* (numerator) compared to *GBAP1* (denominator) (frontal lobe, log2 fold change = 2.3; hippocampus, log2 fold change = 3.1). That is, quantification with short-read RNA-seq wrongly estimated the relative expression of this parent-pseudogene pair by a 2-3 log2 fold difference (frontal cortex, Grubbs’ test statistic = 3.58, *P* = 0.03; hippocampus, Grubbs’ test statistic = 4.27, *P* < 0.01, Grubbs test for one outlier) (**Fig. 3c**).

### Long-read RNA-sequencing reveals novel potentially protein-coding transcripts for *GBA1* and *GBAP1* with no dominant transcript in the human brain

The inaccuracies in quantification suggested that high dependence on short-read RNA-seq technologies may have also led to inaccuracies in *GBA1* and *GBAP1* transcript structures. To address this, we performed targeted Pacific Biosciences (PacBio) isoform sequencing (Iso-Seq) (**Extended Data Fig. 2a**) on 12 human brain regions. Brain tissue was used because of *GBA1*’s importance in neurological disease^12–15,35,36^, and previous evidence suggesting that transcriptome annotation is most incomplete in human brain^37^. We used PacBio Iso-Seq, which has >99% base pair accuracy enabled by circular consensus sequencing (CCS), which in turn, allows accurate mapping. To ensure that full-length reads were generated from mature mRNA alone, we used high-quality polyadenylated RNA (RNA integrity number > 8) pooled from multiple individuals per tissue (**Supplementary Table 2**). *GBA1* and *GBAP1* cDNAs were enriched using biotinylated hybridization probes designed against exonic and intronic genic regions (**Supplementary fig. 4**) to ensure that few assumptions were made regarding transcript structure. Collapsing mapped reads resulted in 2,368 *GBA1* and 3,083 *GBAP1* unique transcripts, each supported by ≥ 2 full-length (FL) HiFi reads across all samples (**Extended Data Fig. 3a,b**). After QC (Quality Control) and filtering for a minimum of 0.3% transcript usage per sample (equating to a mean of 43.4 – 11,127.2 and 15.4 – 1,161.3 FL HiFi reads for *GBA1* and *GBAP1* respectively) we identified 32 *GBA1* and 48 *GBAP1* transcripts (**Fig. 4**), thus providing the most reliable annotation of *GBA1* and *GBAP1* transcription to date.

**Fig. 4:**
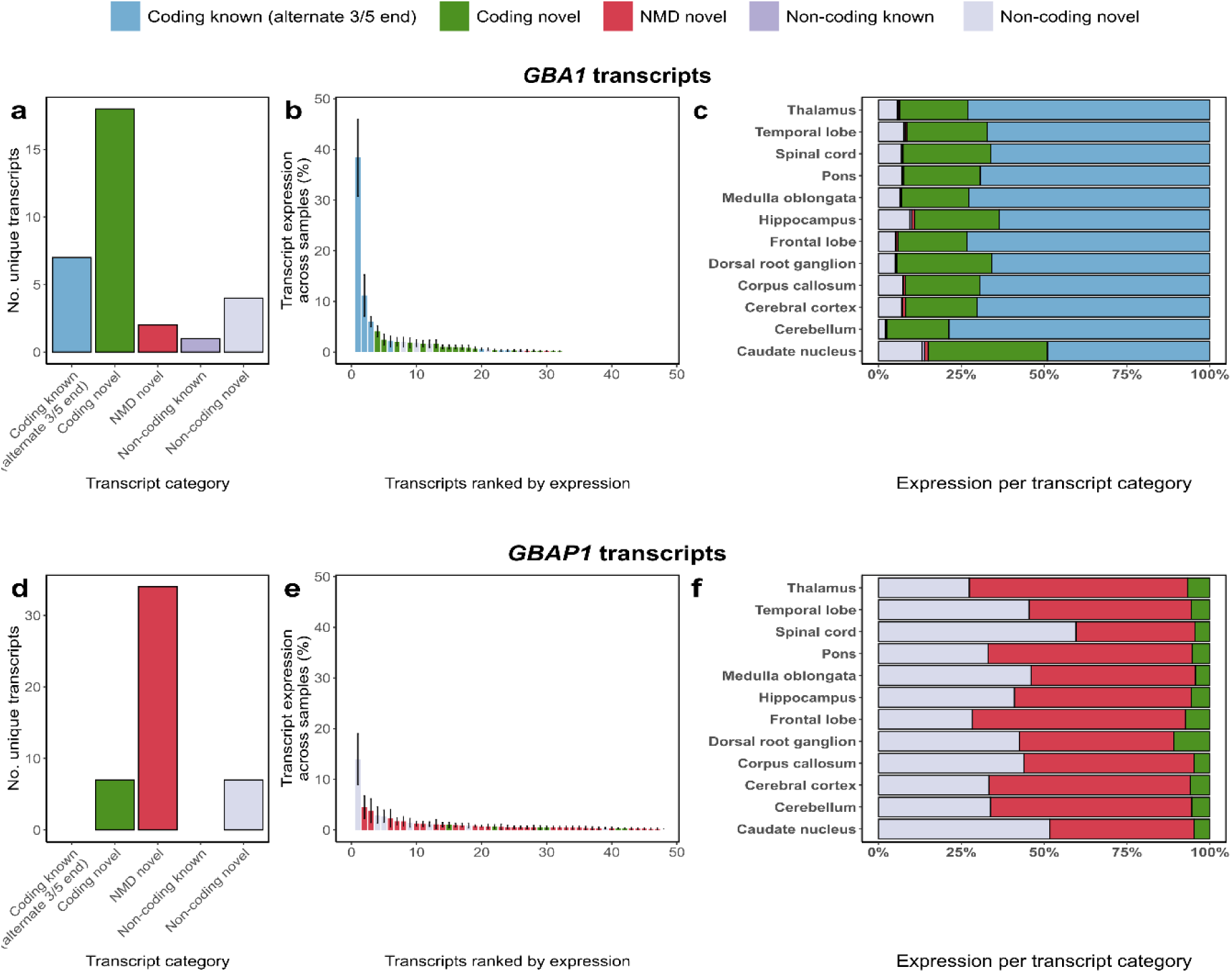
Targeted long-read RNA-sequencing of *GBA1* and *GBAP1* identifies frequent novel transcription. **a,** Bar chart depicting the number of unique *GBA1* transcripts identified per transcript category through targeted long-read RNA-sequencing across 12 human brain regions. **b,** Normalised expression per *GBA1* transcripts corresponding to the percentage of expression per transcripts out of total expression of the loci. **c,** Stacked bar chart showing expression per transcript category of *GBA1* across 12 human brain regions. **d,** Bar chart depicting the number of unique *GBAP1* transcripts identified per transcript category through targeted long-read RNA-sequencing across 12 human brain regions. **e,** Normalised expression per *GBAP1* transcripts corresponding to the percentage of expression per transcripts out of total expression of the loci. **f,** Stacked bar chart showing the expression per transcript category of *GBAP1* across 12 human brain regions.

Next, we examined the identified transcripts for coding potential, nonsense-mediated decay (NMD) and similarity with the existing annotation from GENCODE, to categorise transcripts into the following five categories: (1) coding known (alternate 3’/5’ end); (2) coding novel; (3) NMD novel; (4) non-coding known; and (5) non-coding novel (**Methods** and **Extended Data Fig. 2b**). We noted that 24 of the 32 identified *GBA1* transcripts and all 48 identified *GBAP1* transcripts were absent from GENCODE (**Fig. 4a,d**).

Contrary to the expectation that most protein-coding genes express one dominant transcript^38–40^, we did not find a dominant *GBA1* or *GBAP1* transcript across any of the 12 brain regions sequenced. In fact, the most highly expressed *GBA1* transcript (PB.845.2786; a full splice match to ENST00000368373), only corresponded to a mean of 38.4 ± 7.6% of total transcription at the locus (**Fig. 4b**). Although less surprising for a pseudogene, the most highly expressed transcript of *GBAP1* (Non-coding novel) only corresponded to a mean of 14.0 ± 5.0% of total transcription at the locus (**Fig. 4e**).

### Collectively 25 novel protein-coding transcripts of *GBA1* and its pseudogene *GBAP1* are identified

We found that of all the coding transcripts detected, 18 *GBA1* transcripts had a novel open reading frame (ORF) and 7 *GBAP1* transcripts were predicted to encode a protein, despite *GBAP1* being classified as a pseudogene (**Fig. 4a,d**). Since usage of unannotated 5’ transcription start sites (TSSs) was a common feature of *GBA1* and *GBAP1* transcripts with novel ORFs (open reading frames) (**Supplementary Fig. 5**), we focused on validating these sites using Cap Analysis Gene Expression (CAGE) peaks (defined by FANTOM5^41,42^). We found, despite the fact that CAGE seq only captures the first 20–30 nucleotides from the 5’-end (unique mapping only), 57% (*n* = 4) and 50% (*n* = 9) of novel *GBA1* and *GBAP1* 5’ TSSs, respectively, were located within 50 bp of CAGE peaks providing additional confidence in calling of these transcripts. Moreover, we validated all novel ORFs through additional targeted Iso-Seq of *GBA1* and *GBAP1* in iPSC-derived cortical neurons (*n* = 6), astrocytes (*n* = 3), and microglia (*n* = 3). In summary, we were able to detect *GBA1* and *GBAP1* transcripts with novel ORFs using a different RNA-seq technology and validate them in an independent data set.

To explore the coding potential of *GBA1* and *GBAP1* transcripts with novel ORFs, we employed a sequence-based approach along with AlphaFold2^43^ (which accurately predicts GBA1 structure; **Supplementary Fig. 6**). We focused on the most highly expressed *GBA1* (*n* = 3) and *GBAP1* (*n* = 2) ORFs (**Fig. 5a, b**). Although protein isoforms of both genes were predicted to have highly similar tertiary structures at the C-terminus, we predicted that all protein products would be unlikely to have GCase activity due to the partial/full loss of key enzymatic sites, or the absence of the lysosomal targeting sequence (LIMP2-interface region; **Fig. 5c-h** and **Extended Data Fig. 4**)^44,45^. To assess the coding potential of these novel *GBA1* and *GBAP1* transcripts, we amplified the ORFs and cloned them into a vector with a C-terminus FLAG tag. We transfected these vectors into H4 cells with homozygous knockout of *GBA1*, and found translation of all transcripts as detected with both an anti-FLAG antibody and an antibody directed to the conserved C-terminus (**Fig. 6a** and **Supplementary Fig. 7**). However, none of these transcripts encoded protein isoforms with GCase activity, including those transcribed from *GBAP1* (**Fig. 6b**). We also found no evidence to suggest that these protein isoforms inhibited constitutive GCase activity in H4 parental cells expressing GBA1 (**Fig. 6c**). Furthermore, immunohistochemical analysis in the H4 GBA1 KO and the H4 parental line (expressing endogenous GBA1) showed the lack of lysosomal localization of novel GBA1 and GBAP1 protein isoforms, as predicted (**Fig. 6d**).

**Fig. 5:**
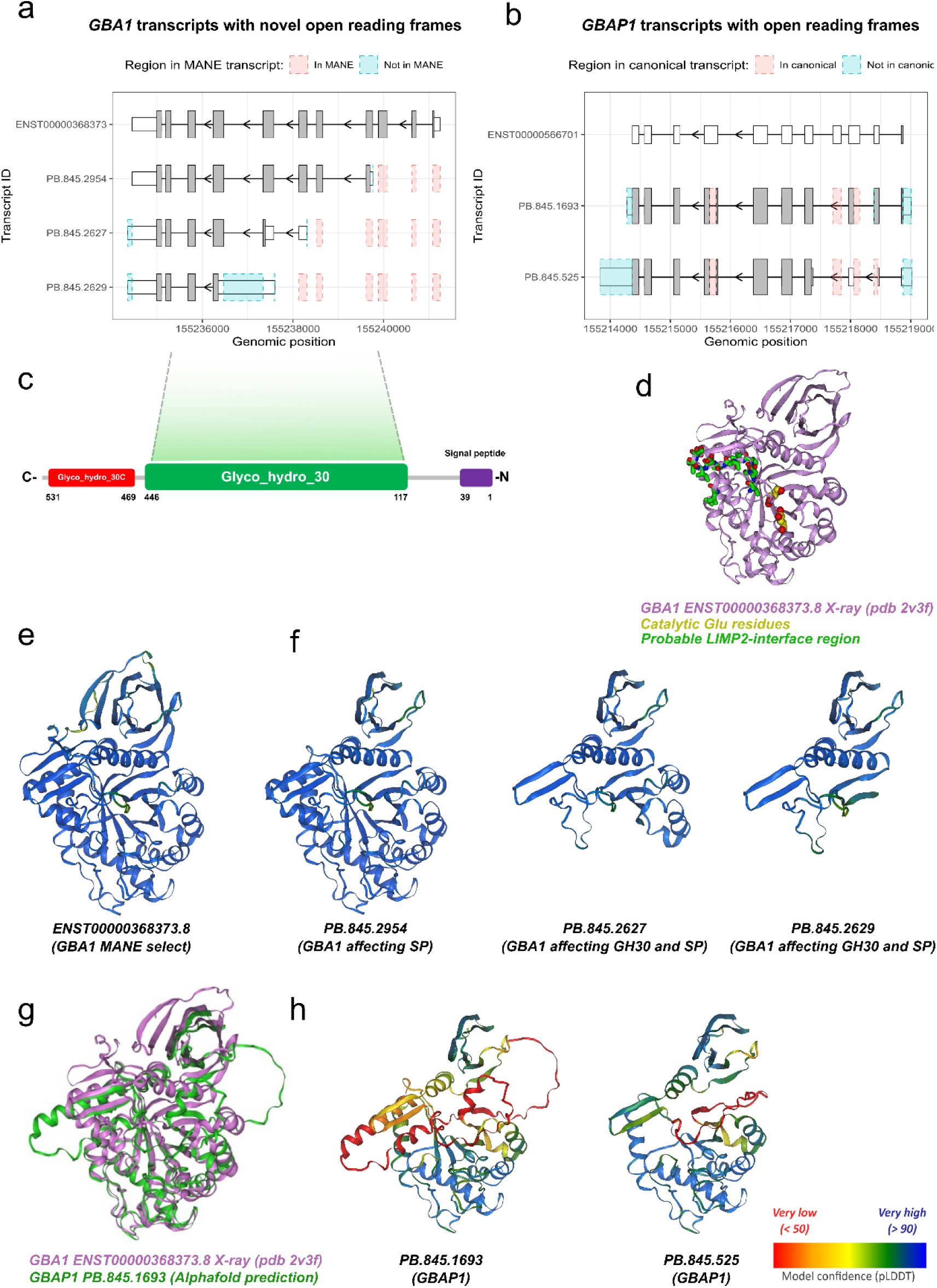
Novel protein-coding transcripts of *GBA1* and *GBAP1* share a similar structure at the C-terminus but with partial or full loss of key domains. **a,** Novel coding *GBA1* transcripts plotted using ggtranscript with differences as compared to MANE select (ENST00000368373) highlighted in blue and red. **b,** Novel predicted coding *GBAP1* transcripts plotted using ggtranscript with differences as compared to ensemble canonical (ENST00000566701) highlighted in blue and red. **c,** Schematic representation of GBA1 with the signal peptide (amino acids 1-39), glyco_hydro_30 (amino acids 117-446), and glycol_hydro_30C (amino acids 469-531). **d,** X-ray structure of GBA1 (PDB 2v3f) with catalytic Glu residues highlighted in yellow and probable LIMP-2 interface region highlighted in purple. **e,** Alphafold2 predictions of GBA1 MANE select (ENST00000368373) and **f,** the three most highly expressed novel protein-coding GBA1 isoforms colored by prediction confidence score (pLDDT). **g,** X-ray structure of GBA1 (PDB 2v3f) (violet) superimposed on AlphaFold2 predicted structure of the longer ORF generated by *GBAP1* PB.845.1693 (green). **h,** Alphafold2 predictions of the two most highly expressed novel protein-coding GBAP1 isoforms colored by prediction confidence score (pLDDT).

**Fig. 6:**
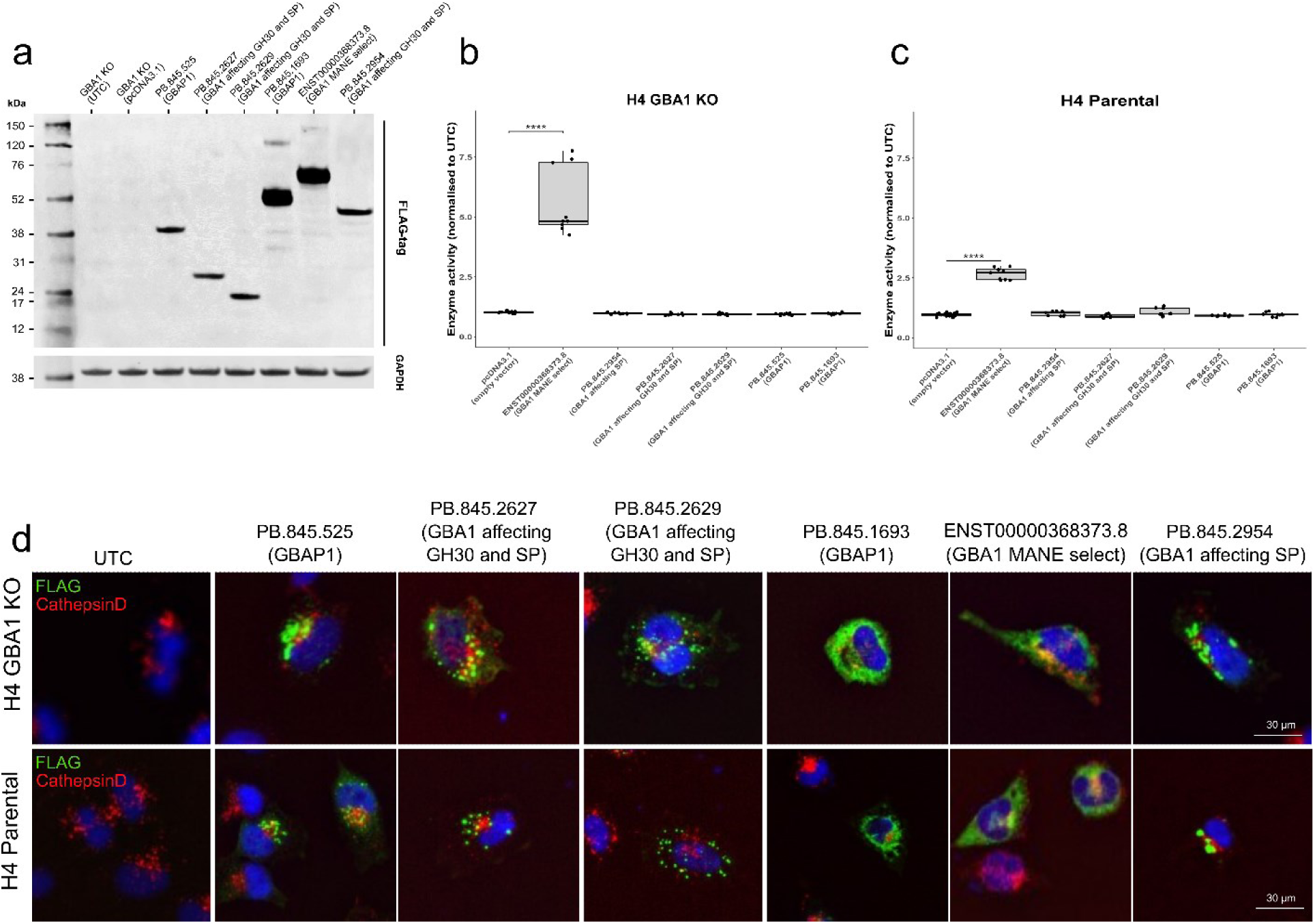
Novel GBA1 and GBAP1 transcripts are translated with no GCase activity and impaired lysosomal co-localization. **a,** Immunoblot of H4 GBA (−/−/−) knockout cells transiently transfected with GBA1 and GBAP1 constructs containing a c-terminus FLAG-tag. GBA1 and GBAP1 expression was detected using FLAG-tag antibody, GAPDH was used as a loading control. The predicted protein sizes are: PB.845.525 (GBAP1; 321 aa; 35 kDa), PB.845.2627 (GBA1 affecting GH30 and SP; 219 aa; 24 kDa), PB.845.2629 (GBA1 affecting GH30 and SP; 164 aa; 18 kDa), PB.845.1693 (GBAP1; 399 aa; 44 kDa), ENST00000368373 (GBA1 MANE select; 537 aa; 62 kDa) and PB.845.2954 (GBA1 affecting GH30 and SP; 414 aa; 46 kDa). **b,** Lysosomal enzyme assay of H4 GBA (−/−/−) knockout cells transiently transfected with GBA1 and GBAP1 constructs, **c,** and in H4 parental. GCase enzyme activity was significantly increased only in H4 parental and GBA (−/−/−) knockout cells transiently transfected with the GBA1 full-length construct (ENST00000368373), compared to the empty vector control (n=3). **d,** Lysosomal co-localisation is impaired in novel GBA1 and GBAP1 transcripts. Immunohistochemistry of H4 parental and GBA (−/−/−) knockout cells transiently transfected with GBA1 and GBAP1 constructs containing a c-terminus Flag tag. Co-localisation of GBA-Flag and GBAP1-Flag (Green) with CathepsinD (Red) was detected using Flag tag antibody.

To explore translation *in vivo*, we interrogated public untargeted mass spectrometry data of human prefrontal cortex^46^. Since novel GBA1 isoforms have no unique sequences that differentiate them, we focused on GBAP1 isoforms. We found proteomic support for GBAP1 (PB.845.1693) within the dataset with a protein Q-value of <0.01. In particular, we identified the unique amino acid sequence QWALDGAEYR, which is unique to GBAP1 and was not identified when searched within the UniProt human protein reviewed dataset. This shows translation of GBAP1 within the human prefrontal cortex.

### *GBA1* and *GBAP1* transcripts show cell-type-selectivity in human brain

We found that novel protein-coding transcripts of *GBA1* without predicted GCase activity were common, collectively accounting for between 15.8% (cerebellum) −31.7% (caudate nucleus) of transcription from the *GBA1* locus. Notably, we found that only 48% of transcription in the caudate nucleus was predicted to encode a protein isoform with GCase activity. This high variability in the usage of *GBA1* transcripts with novel ORFs across the human brain led us to hypothesise that these transcripts may have high cell type specificity. To test this, we employed both 5’ single-nucleus RNA-seq (snRNA-seq) of human dorsolateral prefrontal cortex (DLPFC) and targeted PacBio Iso-Seq of human iPSC-derived brain-relevant cell types. Our analysis revealed cell-type-specific differences in the expression of *GBA1* and *GBAP1* (**Fig. 7**).

**Fig. 7:**
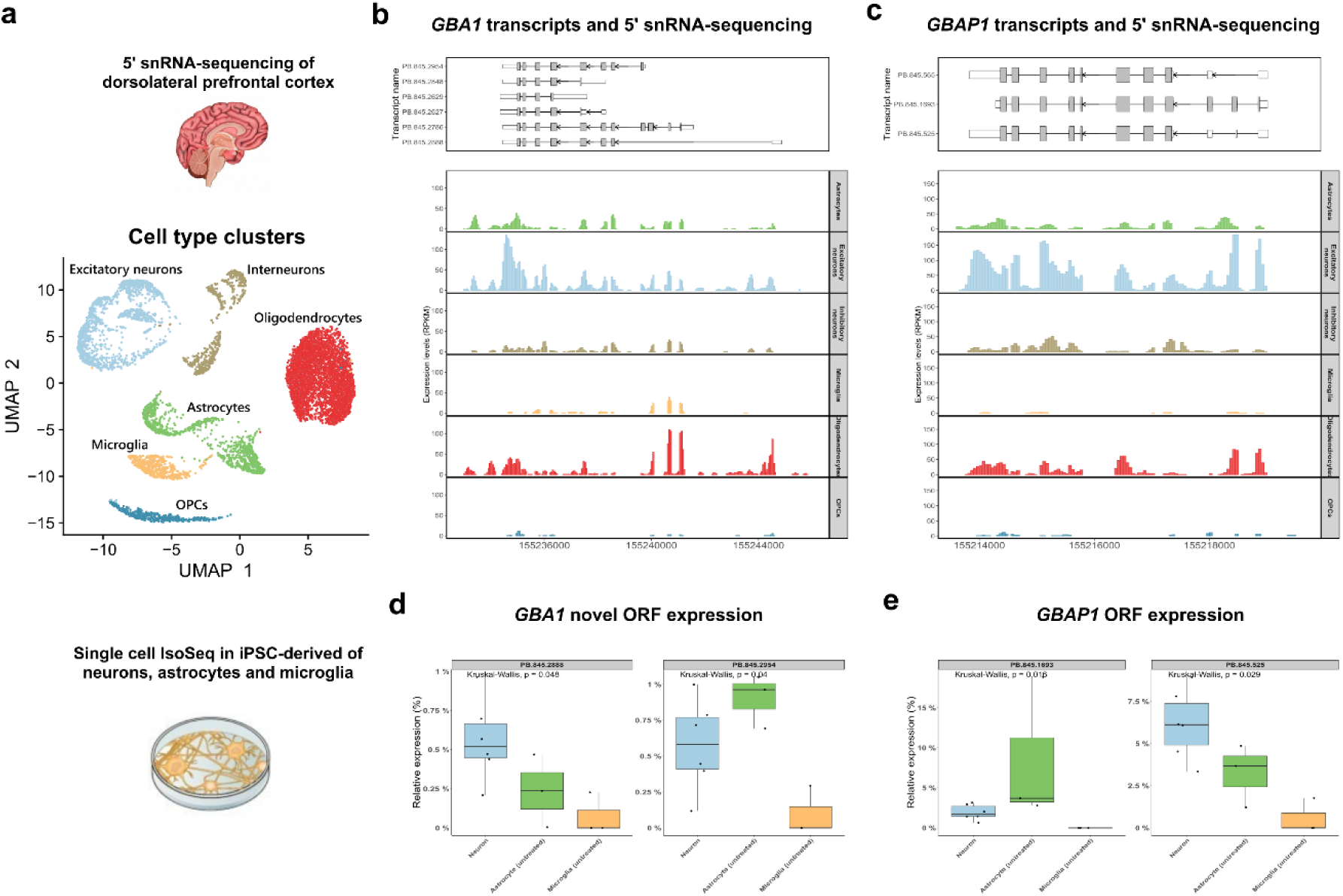
Novel protein-coding transcripts of *GBA1* and *GBAP1* shows cell type specific usage. **a,** UMAP labelled by characterized cell types in human dorsolateral prefrontal cortex. **b,** *GBA1* expression from 5’ single-nucleus RNA-sequencing of human dorsolateral prefrontal cortex. **c,** *GBAP1* expression from 5’ single-nucleus RNA-sequencing of human dorsolateral prefrontal cortex. **d,** Expression of *GBA1* ORFs from PacBio Iso-Seq data generated from human iPSC-derived cortical neuron (n = 6), astrocyte (n = 3) and microglia (n = 3) cultures. **e,** Expression of *GBAP1* ORFs from PacBio Iso-Seq data generated from human iPSC-derived cortical neuron (n = 6), astrocyte (n = 3) and microglia (n = 3) cultures.

Specifically, we used 5’ snRNA-seq of DLPFC to assess the expression of *GBA1* and *GBAP1* in various cell types, including astrocytes, excitatory neurons, inhibitory neurons, microglia, oligodendrocytes, and oligodendrocyte precursor cells (OPCs) (**Fig. 7a**). Our analysis showed an absence of signal at the first exon of PB.845.2888 (*GBA1*) in microglia, along with an overall lower expression of novel *GBA1* transcripts in microglia and OPCs (Fig. 7b). Interestingly, we found that microglia showed significantly lower relative expression of shorter *GBA1* ORFs lacking GCase activity (PB.275.2954 and PB.845.2888) compared to neurons or astrocytes, using PacBio Iso-Seq of human iPSC-derived neurons, astrocytes, and microglia (**Fig. 7d**).

Likewise, our analysis revealed that excitatory neurons had higher expression of *GBAP1* ORF transcripts as compared to microglia, using 5’ snRNA-seq of DLPFC (**Fig. 7c**). Further, using PacBio Iso-Seq of human iPSC-derived neurons, astrocytes, and microglia, we found significant cell-type-specific differences in *GBAP1* ORF usage, with lower utilization of all *GBAP1* ORFs in microglia compared to excitatory neurons and astrocytes (**Fig. 7e**). Additionally, our profiling of H3K4me3 mark in neurons using CUT&RUN^47^ supported transcriptional activity at the 5’ transcription start sites (TSS) of *GBAP1* ORF transcripts (**Extended Data Fig. 5**).

### Inaccurate annotation is frequent among parent genes across human tissues

We have shown significant inaccuracies in annotation of the parent gene *GBA1*. However, we wanted to explore the scope of this problem. To do so we compared inaccuracies in annotation of all 3,665 parent genes compared with other protein-coding genes (including paralogs). Initially, we used public long-read RNA-seq data from 29 samples (*n*, Brain = 9, Heart = 16 and lung = 6; **Supplementary Table 3**) to assess the proportion of transcripts per gene, with at least one novel splice site in the coding sequence that would result in a novel ORF. Despite a low sequencing depth (mean, 2.2 ± 0.9 million full-length reads per sample), we found a significant increase in such events among parent genes compared to other protein-coding genes (parent genes = 23.9 ± 11.5%; protein-coding genes = 22.7 ± 11.4%; two-sided Wilcoxon rank-sum test *p* < 0.01; **Fig. 8a**). We extended this analysis to a greater number of samples (*n* = 7,595) and human tissues (n = 41, GTEx) using annotation-agnostic short-read RNA-seq analyses to quantify the proportion of parent genes with evidence of novel splicing (**Method**). Based on the identification of novel expressed genomic regions^37^ and novel splice site usage, we found that the proportion of genes with incomplete annotation was significantly higher among parent genes compared to other protein-coding genes (novel expression regions: parent genes = 13.9 ± 1.4%; protein-coding genes = 10.8 ± 1.3%; two-sided Wilcoxon rank-sum test *p* < 0.01; **Fig. 8b**; splice site usage: parent genes = 66.5 ± 3.5%; protein-coding genes = 54.8 ± 4.3; two-sided Wilcoxon rank-sum test *p* < 0.01; **Fig. 8c**). This observation was consistent across all tissues analysed (**Supplementary Fig. 8**).

**Fig. 8:**
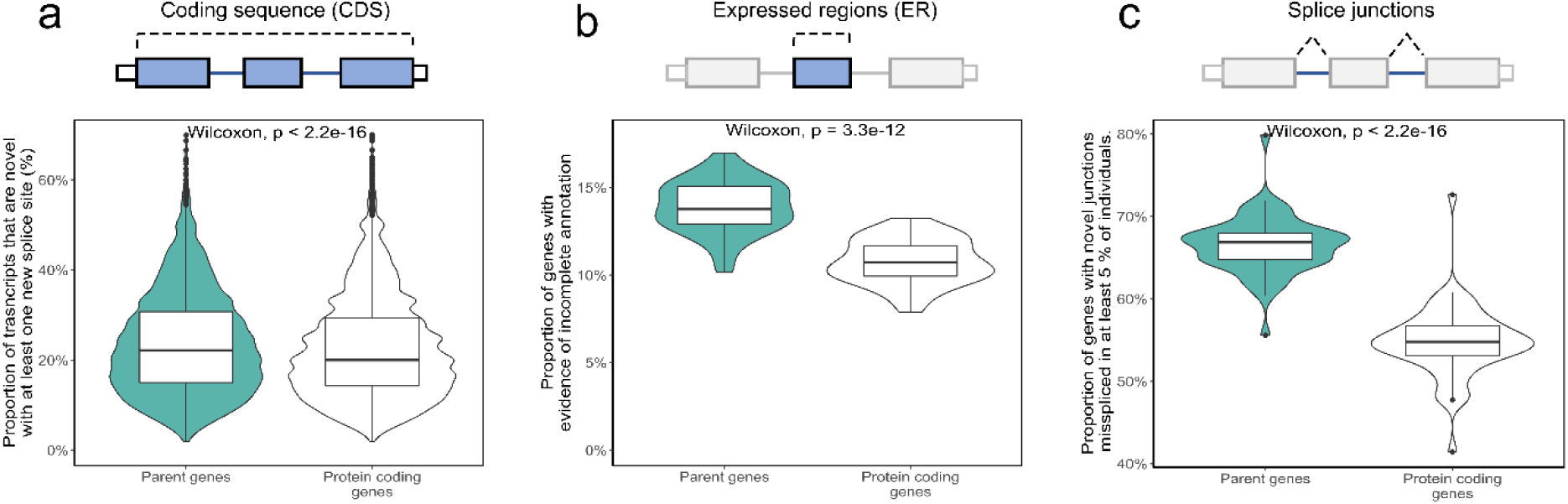
Inaccuracies in annotation is common for parent genes on a genome-wide scale. **a,** Proportion of transcripts per parent gene and per protein-coding gene without a pseudogene with a novel splice site from long-read RNA-sequencing data of 9 frontal cortex samples. **b,** Proportion of genes with evidence of incomplete annotation based on the identification of novel expressed genomic regions from short-read RNA-sequencing data. **c,** Proportion of genes with evidence of incomplete annotation based on the identification novel splice junctions found in at least 5% of samples from short-read RNA-sequencing data.

## DISCUSSION

Here we show that widespread expression and alternative splicing of pseudogenes in human tissues has limited our understanding of both pseudogene and parent gene transcription with a significant impact on our appreciation of gene function. Our long-read RNA-seq analysis of the parent gene *GBA1* and its pseudogene *GBAP1* demonstrated significant diversity in transcription and showed that contrary to expectation ^48,49^, no single transcript dominated expression of either gene in human brain. A substantial portion of transcription from both loci was novel, leading to the identification of novel protein-coding transcripts with tissue and cell-type specific biases in usage. Together these findings have a significant impact on our understanding of the potential mechanisms through which genetic variation at the *GBA1-GBAP1* locus could explain phenotypic diversity in Gaucher disease and modulate disease risk and expressivity in Parkinson’s disease.

Even though current annotation is known to be incomplete, especially in the brain^37^, the extent of transcriptional variety and novelty at parent gene loci was surprising, and particularly so at *GBA1*. After all, GCase dysfunction has been implicated in human disease since 1965^6^ and mutations in *GBA1* have been described since 1987^50^, making *GBA1* one of the most studied genes in the genome. Nonetheless, we found that as much as 31.7% of *GBA1* transcription in the caudate nucleus, a key component of the basal ganglia, may be translated into novel protein isoforms that do not localise in lysosomes and thus, lack GCase activity. Given that reduced GCase activity has been linked to PD^12–15^ and basal ganglia circuitry dysfunction is a feature of this disease, these findings were all the more striking.

We also demonstrate that inaccuracies in annotation were significantly more common across parent genes as compared to other protein coding genes (**Fig. 8**) and were not restricted to *GBA1*. High sequence similarity within the genome and subsequent multimapping of short RNA-sequencing reads, has impacted on our understanding of many genes, including those already causally linked to disease. Such loci are predictable using sequence similarity analyses, the technology to resolve these “problem” loci is available and the impact on our understanding of disease is likely to be significant. As exemplified by *GBA1*-*GBAP1*, our limited understanding of transcription from this locus results in errors in quantification of gene expression and all dependent analysis from differential gene expression in disease to quantitative trait loci detection. Beyond a research setting, inaccuracies in annotation will affect variant interpretation and consequently diagnostic yield for some disease-associated genes. Finally, and perhaps most importantly, significant inaccuracies in transcript annotation impact on our understanding of gene function. Directed by our long-read RNA-sequencing results, we have found that some *GBAP1* transcripts are more highly expressed in neurons and astrocytes, share a similar predicted 3D protein structure to GBA1, have protein products that do not localise to lysosomes and lack GCase activity. Yet, we find robust evidence of translation of such *GBAP1* transcripts in human brain using high-throughput mass spectrometry data^46^. Extrapolating these findings to GBA1, where mass spectrometry data was uninformative, would suggest a non-lysosomal function for both GBA1 and GBAP1 in brain, and particularly in neurons.

We propose that improving our understanding of the molecular functions of parent-pseudogene pairs will become increasingly important to the development and success of RNA-targeting therapies. Accurate annotation is required at the tissue- and cell-level to design effective Antisense Oligonucleotides (ASOs) or gene therapies. Furthermore, some pseudogenes may represent particularly high value therapeutic targets due to their potential to operate as genetic modifiers of Mendelian disorders. In fact, Nusinersen, which targets the splicing of former pseudogene *SMN2*, is a highly successful treatment for Spinal muscular atrophy ^51^. Thus, a deeper understanding of pseudogene function could lead to new and innovative therapeutic strategies.

Taken together, our findings from the *GBA1-GBAP1* study demonstrate the need for thorough re-examination of transcription in duplicated genomic regions, such as parent-pseudogene pairs. By employing accurate full-length transcript sequencing, we are able to resolve these complex loci with unprecedented detail, leading to novel transcript discovery and, as a result, new insights into the functionality of human diseases.

## ONLINE METHODS

### PSEUDOGENES AND PARENTAL GENES

#### Pseudogene and parent gene annotations

Pseudogene annotations were obtained from GENCODE v 38^24^ (https://ftp.ebi.ac.uk/pub/databases/gencode/Gencode_human/). We included all HAVANA annotated pseudogenes excluding polymorphic pseudogenes. Biotypes were clustered using the “gene_type” column so that “IG_V_pseudogene”, “IG_C_pseudogene”, “IG_J_pseudogene”, “IG_pseudogene”, “TR”, “TR_J_pseudogene”, “TR_V_pseudogene”, “transcribed_unitary_pseudogene”, “unitary_pseudogene” = “Unitary”; “rRNA_pseudogene”, “pseudogene” = “Other”; “transcribed_unprocessed_pseudogene”, “unprocessed_pseudogene”, “translated_unprocessed_pseudogene” = “Unprocessed”; “processed_pseudogene”, “transcribed_processed_pseudogene”, “translated_processed_pseudogene” = “Processed”. Parent genes have previously been inferred^25^ and were obtained from psiCube (http://pseudogene.org/psicube/index.html).

#### Expression analysis from GTEx

Pseudogene and parent gene expression was assessed using median transcript per million (TPM) expression per tissue generated by the Genotype-Tissue Expression Consortium (GTEx, v8, accessed on 10/11/2021). As GTEx only use uniquely mapped reads for expression and multimapping was a concern, expression was assessed as a binary variable. That is, a gene with a median TPM > 0 was considered to be expressed.

For quantitative expression of *GBA1* and *GBAP1* we used RNA-seq data for 17,510 human samples originating from 54 different human tissues (GTEx, v8) that was downloaded using the R package recount (v 1.4.6)^52^. Cell lines, sex-specific tissues, and tissues with 10 samples or below were removed. Samples with large chromosomal deletions and duplications or large copy number variation previously associated with disease were filtered out (smafrze != “EXCLUDE”). For any log_2_ fold change calculations *GBA1* is the numerator and *GBAP1* is the denominator.

#### Alternative splicing analysis using long-read RNA-sequencing

To identify alternative splicing of pseudogenes we used publicly available long-read RNA-seq data from ENCODE^53^ (https://www.encodeproject.org/rna-seq/long-read-rna-seq/). We included 29 samples from Brain (*n* = 9), Heart (*n* = 16) and lung (*n* = 6). A description of the samples included can be found in **Supplementary Table 2**. All samples were sequenced on the PacBio Sequel II platform and processed with the ENCODE DCC deployment of the TALON pipeline (v2.0.0; https://github.com/ENCODE-DCC/long-read-rna-pipeline)^54^.

#### Online Mendelian Inheritance in Man data

Phenotype relationships and clinical synopses of all Online Mendelian Inheritance in Man (OMIM) genes were downloaded using API through https://omim.org/api (accessed 14/04/2022)^26^. Parent genes were annotated genes as OMIM morbid if they were listed as causing a mendelian phenotype.

#### Sequence similarity

Sequence similarity of parent genes and pseudogenes has previously been calculated by Pei *et al*.^2^ and is available through The Pseudogene Decoration Resource (psiDr; http://www.pseudogene.org/psidr/similarity.dat; accessed 14/04/2022). We compared the sequence similarity of parent and pseudogenes considering the coding sequence (CDS) of parent genes.

#### Multimapping from short-read RNA-sequencing

Multimapping rates of parent genes, including *GBA1* and *GBAP1*, were investigated in human anterior cingulate cortex samples previously reported in Feleke & Reynolds et al^34^. Here, we used control individuals (*n* = 5) and individuals with Parkinson’s disease (PD) with or without dementia (*n* = 13). Adapter trimming and read quality filtering was performed with default options using Fastp (v 0.23.2; RRID:SCR_016962)^55^, with quality control metrics generated using both Fastp and FastQC (v 0.11.9; RRID:SCR_014583). Alignment to the GRCh38 genome using GENCODE v 38 was performed using STAR (v 2.7.10; RRID:SCR_004463)^56^. ENCODE standard options for long RNA-seq were used with STAR, except for alignSJDBoverhangMin, outSAMmultNmax and outFilterMultimapNmax. outFilterMultimapNmax sets the rate of multimapping permitted; as a conservative estimate we set this to 10, half the ENCODE standard. outSAMmultNmax was set to −1, which allowed multimapped reads to be kept in the same output SAM/BAM file. The QC and alignment processes were performed using a nextflow^57^ pipeline. BAM files were sorted and indexed using Samtools (v 1.14; RRID:SCR_002105) ^58^ and filtered in R (v 4.0.5; RRID:SCR_001905) for reads overlapping the *GBA1* or *GBAP1* locus, using GenomicRanges (v 1.42.0; RRID:SCR_000025)^59^ and Rsamtools (version 2.6.0). Only paired first mate reads on the correct strand (minus for both *GBA1* and *GBAP1*) selected. The “NH” tag, which provides the number of alignments for a read was also extracted from the SAM header. The CIGAR string of the read was used to provide a width of the reads relative to the reference by adding operations that consume the reference together. Reads were then filtered, using dplyr (v 1.0.9; RRID:SCR_016708)^60^ and tibble (v 3.1.6)^60^, with this new width to leave reads that aligned completely within the *GBA1* and *GBAP1* locus. Reads were then split between unique alignment and multimapping alignments based on the “NH” tag. The percentage of reads (uniquely mapped / (uniquely mapped + multimapped)) that mapped uniquely to either the *GBA1* or *GBAP1* locus was then calculated. Additionally, for reads that multimapped to the *GBA1* or *GBAP1* locus the read name was extracted and searched for within the reads that multimapped to the alternate locus (i.e., reads names from reads that multimapped to the *GBA1* locus were searched against read names for reads that multimapped to the *GBAP1* locus). This provided a percentage of reads that aligned to *GBA1* that that also aligned elsewhere and the percentage of reads aligning to *GBAP1*. Code and commentary can be found here: https://github.com/Jbrenton191/GBA_multimapping_2022.

### OXFORD NANOPORE DIRECT CDNA SEQUENCING

#### Samples

Human Poly A+ RNA of healthy individuals that passed away from sudden death/trauma derived from frontal lobe and hippocampus were commercially purchased through Clontech (**Supplementary Table 2**).

#### Direct cDNA sequencing

A total of 100ng of Poly A+ RNA per sample was used for initial cDNA synthesis and subsequent library preparation according to the direct cDNA sequencing (SQK-DCS109) protocol described in detail at protocols.io (dx.doi.org/10.17504/protocols.io.yxmvmkpxng3p/v1). Sequencing was performed on the PromethION using one R9.4.1 flow cell per sample and base-called using Guppy (v 4.0.11; Oxford Nanopore Technologies—ONT, Oxford, UK). Resulting fastq files were processed through the “pipeline-nanopore-ref-isoforms” (https://github.com/nanoporetech/pipeline-nanopore-ref-isoforms). Gene abundances was calculated implementing the −A parameter in StringTie (v 2.1.1 RRID:SCR_016323)^61^. Data is available and deposited in the Gene Expression Omnibus under accession GSE215459

#### Comparing short-read quantification versus long-read quantification

For each sample in GTEx a log2 fold change was calculated with *GBA1* as the numerator and *GBAP1* as the denominator across frontal lobe and hippocampus. Shapiro-Wilk normality test in each tissue was used to confirm a normal distribution. To compare against ONT long-read quantification we used Grubbs’ test (maximum normalized residual test) for a single outlier.

### PACBIO TARGETED ISO-SEQ

#### Samples

##### Human brain samples

Human Poly A+ RNA of healthy individuals that passed away from sudden death/trauma derived from caudate nucleus, cerebellum, cerebral cortex, corpus callosum, dorsal root ganglion, frontal lobe, hippocampus, medulla oblongata, pons, spinal cord, temporal lobe, and thalamus were commercially purchased through Clontech (**Supplementary Table 2**).

##### iPSC, neuroepithelial, neural progenitor, cortical neuron, astrocyte, and microglia cells

Control iPSCs consisted of the previously characterized lines Ctrl1^62^, ND41866 (Coriel), RBi001 (EBiSC/Sigma) and SIGi1001 (EBiSC/Sigma) as well as the isogenic line previously generated^63^. Reagents were purchased from Thermo Fisher Scientific unless otherwise stated. iPSCs lines were grown in Essential 8 media on geltrex substrate and passaged using 0.5M EDTA. Cortical neurons were differentiated using dual SMAD inhibition for 10 days (10µM SB431542 and 1µM dorsomorphin, Tocris) in N2B27 media before maturation in N2B27 alone^64^. Day 100 +/- 5 days was taken as the final timepoint. Astrocytes were generated following a similar neural induction protocol until day 80 before repeatedly passaging cortical neuronal inductions in 10ng/ml FGF2 (Peprotech) to enrich for astrocyte precursors. At day 150, to generate mature astrocytes, a two-week maturation consisted of BMP4 (10ng/ml, Thermo Fisher) and LIF (10ng/ml, Sigma)^65^. To induce inflammatory conditions, astrocytes were stimulated with TNFα (30ng/ml, Peprotech), IL1α (3ng/ml, Peprotech) and C1q (400ng/ml, Merck)^66^. iPSC-microglia were differentiated following the protocol of Xiang at al^67^. Embryoid bodies were generated using 10,000 iPSCs and myeloid differentiation was initiated in Lonza XVivo 15 media, IL3 (25ng/ml, Peprotech) and MCSF (100ng/ml, Peprotech). Microglia released from embryoid bodies were harvested weekly from 4 weeks and matured in DMEM-F12 supplemented with 2% insulin/transferrin/selenium, 1% N2 supplement, 1X glutamax, 1X NEAA and 5ng/ml insulin supplemented with IL34 (100ng/ml, Peprotech), MCSF (25ng/ml, Peprotech), TGFβ1 (5ng/ml, Peprotech). A final two-day maturation consisted of CXC3L1 (100ng/ml, Peprotech) and CD200 (100ng/ml, 2B Scientific). Inflammation was stimulated with lipopolysaccharide (10ng/ml, Sigma). Total RNA was extracted using the Qiagen RNeasy kit according to the manufacturer’s protocol with β-mercaptoethanol added to buffer RLT and with a DNase digestion step included.

#### cDNA synthesis

A total of 250ng of RNA was used per sample for reverse transcription. Two different cDNA synthesis approaches were used: (i) Human brain cDNA was generated by SMARTer PCR cDNA synthesis (Takara) and (ii) iPSC derived cell lines were generated using NEBNext® Single Cell/Low Input cDNA Synthesis & Amplification Module (New England Biolabs). For both reactions sample-specific barcoded oligo dT (12 µM) with PacBio 16mer barcode sequences were added (**Supplementary Table 3**).

##### SMARTer PCR cDNA synthesis

First strand synthesis was performed as per manufacturer instructions, using sample-specific barcoded primers instead of the 3’ SMART CDS Primer II A. We used a 90 min incubation to generate full-length cDNAs. cDNA amplification was performed using a single primer (5’ PCR Primer II A from the SMARTer kit, 5′ AAG CAG TGG TAT CAA CGC AGA GTA C 3′) and was used for all PCR reactions post reverse transcription. We followed the manufacturer’s protocol with our determined optimal number of 18 cycles for amplification; this was used for all samples. We used a 6 min extension time in order to capture longer cDNA transcripts. PCR products were purified separately with 1X ProNex® Beads.

##### NEBNext® Single Cell/Low Input cDNA Synthesis & Amplification Module

A reaction mix of 5.4 μL of total RNA (250 ng in total), 2 μL of barcoded primer, 1.6 μL of dNTP (25 mM) held at 70°C for 5 min. This reaction mix was then combined with 5 μL of NEBNext Single Cell RT Buffer, 3 μL of nuclease-free H_2_O and 2 μL NEBNext Single Cell RT Enzyme Mix. The reverse transcription mix was then placed in a thermocycler at 42°C with the lid at 52°C for 75 minutes then held at 4°C. On ice, we added 1 μL of Iso-Seq Express Template Switching Oligo and then placed the reaction mix in a thermocycler at 42°C with the lid at 52°C for 15 minutes. We then added 30 μL elution buffer (EB) to the 20 μL Reverse Transcription and Template Switching reaction (for a total of 50 μL), which was then purified with 1X ProNex® Beads and eluted in 46 μL of EB. cDNA amplification was performed by combining the eluted Reverse Transcription and Template Switching reaction with 50 μL of NEBNext Single Cell cDNA PCR Master Mix, 2 μL of NEBNext Single Cell cDNA PCR Primer, 2 μL of Iso-Seq Express cDNA PCR Primer and 0.5 μL of NEBNext Cell Lysis Buffer.

#### cDNA Capture Using IDT Xgen® Lockdown® Probes

We used the xGen Hyb Panel Design Tool (https://eu.idtdna.com/site/order/designtool/index/XGENDESIGN) to design non-overlapping 120-mer hybridization probes against *GBA1* and *GBAP1*. We removed any overlapping probes with repetitive sequences (repeatmasker) and to reduce the density of probes mapping to intronic regions 0.2, which means 1 probe per 1.2kb. In the end, our probe pool consisted of 119 probes of which 54 were targeted towards *GBA1* and 65 were targeted towards *GBAP1*.

We pooled an equal mass of barcoded cDNA for a total of 500 ng per capture reaction. Pooled cDNA was combined with 7.5 μL of Cot DNA in a 1.5 mL LoBind tube. We then added 1.8X of ProNex beads to the cDNA pool with Cot DNA, gently mixed the reaction mix 10 times (using a pipette) and incubated for 10 min at room temperature. After two washes with 200 μL of freshly prepared 80% ethanol, we removed any residual ethanol and immediately added 19 μL hybridization mix consisting of: 9.5 μL of 2X Hybridization Buffer, 3 μL of Hybridization Buffer Enhancer, 1 μL of xGen Asym TSO block (25 nmole), 1 μL of polyT block (25 nmole) and 4.5 μL of 1X xGen Lockdown Probe pool.

The PacBio targeted Iso-Seq protocol is described in detail at protocols.io (dx.doi.org/10.17504/protocols.io.n92ld9wy9g5b/v1).

#### Automated Analysis of Iso-Seq data using Snakemake

For the analysis of targeted PacBio Iso-Seq data, we created two Snakemake^68^ (v 5.32.2; RRID:SCR_003475) pipelines to analyse targeted long-read RNA-seq robustly and systematically:

**APTARS** (Analysis of PacBio TARgeted Sequencing, https://github.com/sid-sethi/APTARS): For each SMRT cell, two files were required for processing: (i) a subreads.bam and (ii) a FASTA file with primer sequences, including barcode sequences.

Each sequencing run was processed by ccs (v 5.0.0; RRID:SCR_021174; https://ccs.how/), which combines multiple subreads of the same SMRTbell molecule and to produce one highly accurate consensus sequence, also called a HiFi read (≥ Q20). We used the following parameters: --minLength 10 –maxLength 50000 – minPasses 3 –minSnr 2.5 –maxPoaCoverage 0 –minPredictedAccuracy 0.99. Identification of barcodes, demultiplexing and removal of primers was then performed using lima (v 2.0.0; https://lima.how/) invoking –isoseq –peek-guess. Isoseq3 (v 3.4.0; https://github.com/PacificBiosciences/IsoSeq) was then used to (i) remove polyA tails and (ii) identify and remove concatemers using, with the following parameters refine –require-polya, --log-level DEBUG. This was followed by clustering and polishing with the following parameters using: cluster flnc.fofn clustered.bam – verbose –use-qvs.

Reads with predicted accuracy ≥ 0.99 were aligned to the GRCh38 reference genome using minimap2^69^ (v 2.17; RRID:SCR_018550) using -ax splice:hq -uf –secondary=no. samtools^58^ (RRID:SCR_002105; http://www.htslib.org/) was then used to sort and filter the output SAM for the locus of gene of interest, as defined in the config.yml.

We used cDNA_Cupcake (v 22.0.0; https://github.com/Magdoll/cDNA_Cupcake) to: (i) collapse redundant transcripts, using collapse_isoforms_by_sam.py (--dun-merge-5-shorter) and (ii) obtain read counts per sample, using get_abundance_post_collapse.py followed by demux_isoseq_with_genome.py.

Isoforms detected were characterized and classified using SQANTI3^70^ (v 4.2; https://github.com/ConesaLab/SQANTI3) in combination with GENCODE (v 38) comprehensive gene annotation. An isoform was classified as full splice match (FSM) if it aligned with reference genome with the same splice junctions and contained the same number of exons, incomplete splice match (ISM) if it contained fewer 5′ exons than reference genome, novel in catalog (NIC) if it is a novel isoform containing a combination of known donor or acceptor sites, or novel not in catalog (NNC) if it is a novel isoform with at least one novel donor or acceptor site.

**PSQAN** (Post Sqanti QC Analysis, https://github.com/sid-sethi/PSQAN) Following transcript characterisation from SQANTI3, we applied a set of filtering criteria to remove potential genomic contamination and rare PCR artifacts. We removed an isoform if: (1) the percent of genomic “A” s in the downstream 20 bp window was more than 80% (“perc_A_downstream_TTS” > 80); (2) one of the junctions was predicted to be template switching artifact (“RTS_stage” = TRUE); or (3) it was not associated with the gene of interest. Using SQANTI’s output of ORF prediction, NMD prediction and structural categorisation based on comparison with the reference annotation (GENCODE), we grouped the identified isoforms into the following categories: (1) **Non-coding novel** – if predicted to be non-coding and not a full-splice match with the reference; (2) **Non-coding known** – if predicted to be non-coding and a full-splice match with the reference; (3) **NMD novel** – if predicted to be coding & NMD, and not a full-splice match with the reference; (4) **NMD known** – if predicted to be coding & NMD, and a full-splice match with the reference; (5) **Coding novel** – if predicted to be coding & not NMD, and not a full-splice match with the reference; (6) **Coding known (complete match)** – if predicted to be coding & not NMD, and a full-splice & UTR match with the reference; and (7) **Coding known (alternate 3’/5’ end)** – if predicted to be coding & not NMD, and a full-splice match with the reference but with an alternate 3’ end, 5’ end or both 3’ and 5’ end.

Given a transcript *T* in sample *i* with *FLR* as the number of full-length reads mapped to the transcript *T*, we calculated the normalised full-length reads (*NFLR_Ti_*) as the percentage of total transcription in the sample:

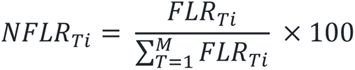

where, *NFLR_Ti_* represents the normalised full-length read count of transcript *T* in sample *i*, *FLR_Ti_* is the full-length read count of transcript *T* in sample *i* and *M* is the total number of transcripts identified to be associated with the gene after filtering. Finally, to summarise the expression of a transcript associated with a gene, we calculated the mean of normalised full-length reads (*NFLR_Ti_*) across all the samples:

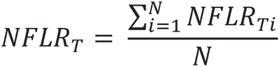

where, *NFLR_T_* represents the mean expression of transcript *T* across all samples and *N* is the total number of samples. To remove low-confidence isoforms arising from artefacts, we only selected isoforms fulfilling the following three criteria: (1) expression of minimum 0.1% of total transcription per sample, i.e., *NFLR_Ti_* ≥ 0.1; (2) a minimum of 80% of total samples passing the *NFLR_Ti_* threshold; and (3) expression of minimum 0.3% of total transcription across samples, i.e., *NFLR_Ti_* ≥ 0.3.

#### Visualizations of transcripts

For any visualization of transcript structures, we have recently developed ggtranscript^71^ (v 0.99.03; https://github.com/dzhang32/ggtranscript), a R package that extends the incredibly popular tool ggplot2^60^ (v 3.3.5 RRID; SCR_014601) for visualizing transcript structure and annotation.

#### CAGE-seq analysis

To assess whether predicted 5’ TSSs of novel transcript were in proximity of Cap Analysis Gene Expression (CAGE) peaks we used data from the FANTOM5 dataset^41,42^. CAGE is based on “cap trapping”: capturing capped full-length RNAs and sequencing only the first 20–30 nucleotides from the 5’-end. CAGE peaks were downloaded from the FANTOM5 project (https://fantom.gsc.riken.jp/5/datafiles/reprocessed/hg38_latest/extra/CAGE_peaks/hg38_liftover+new_CAGE_peaks_phase1and2.bed.gz; accessed 20/05/2022).

### SINGLE NUCLEAR RNA-SEQUENCING

#### Nuclei extraction of cortical post-mortem tissue

Post-mortem brain tissue from control individuals with no known history of neurological or neuropsychiatric symptoms was acquired from the Cambridge Brain Bank (ethical approval from the London-Bloomsbury Research Ethics Committee, REC reference:16/LO/0508). Brains were bisected in the sagittal plane with one half flash-frozen and stored at −80 °C and the other half fixed in 10% neutral buffered formalin for 2–3 weeks. From the flash-frozen blocks, 50-100mg were sampled from the dorsolateral prefrontal cortex (Brodmann area 46) and stored at −80 °C until use. Nuclei were isolated as previously described^72^, with minor modifications^45^. Approximately 20 μg of −80 °C-conserved tissue was thawed and dissociated in ice-cold lysis buffer (0.32M sucrose, 5 mM CaCl2, 3 mM MgAc, 0.1 mM Na2EDTA, 10 mM Tris-HCl pH 8.0, 1 mM DTT) using a 1 mL glass dounce tissue grinder (Wheaton). The homogenate was slowly and carefully layered on top of a sucrose layer (1.8 M sucrose, 3 mM MgAc, 10 mM Tris-HCl pH 8.0, 1 mM DTT) in centrifuge tubes to create a gradient, and then centrifuged at 15,500 rpm for 2 h 15 min. After centrifugation, the supernatant was removed, and the pellet softened for 10 minutes in 100 μL of nuclear storage buffer (15% sucrose, 10 mM Tris-HCl pH 7.2, 70 mM KCl, 2 mM MgCl2) before resuspension in 300 μL of dilution buffer (10 mM Tris-HCl pH 7.2, 70 mM KCl, 2 mM MgCl2, Draq7 1:1000). The suspension was then filtered (70 μm cell strainer) and sorted via FACS (FACS Aria III, BD Biosciences) at 4 °C at a low flowrate, using a 100 μm nozzle (Pipette tips and Eppendorf tubes for transferring nuclei were pre-coated with 1% BSA). 8,500 nuclei were sorted for single-nucleus RNA-seq and then loaded on to the Chromium Next GEM Single Cell 5’ Kit (10x Genomics, PN-1000263). Sequencing libraries were generated with unique dual indices (TT set A) and pooled for sequencing on a NovaSeq 6000 (Illumina) using a 100-cycle kit and 28-10-10-90 reads.

#### Single nucleus RNA-sequencing analysis

Raw base calls were demultiplexed to obtain sample specific FASTQ files using Cell Ranger mkfastq and default parameters (v 6; 10x Genomics; RRID:SCR_017344). Reads were aligned to the GRCh38 genome assembly using the Cell Ranger count (v 6; 10x Genomics; RRID:SCR_017344) with default parameters (--include-introns were used for nuclei mapping)^73^. Nuclei were filtered based on the number of genes detected - nuclei with less of the mean minus a standard deviation, or more than the mean plus two standard deviations were discarded to exclude low quality nuclei or possible doublets. The data was normalized to center log ratio (CLR) to reduce sequencing depth variability. Clusters were defined with Seurat function FindClusters (v; RRID:SCR_007322), using resolution of 0.5. Obtained clusters were manually annotated using canonical marker gene expression as following:

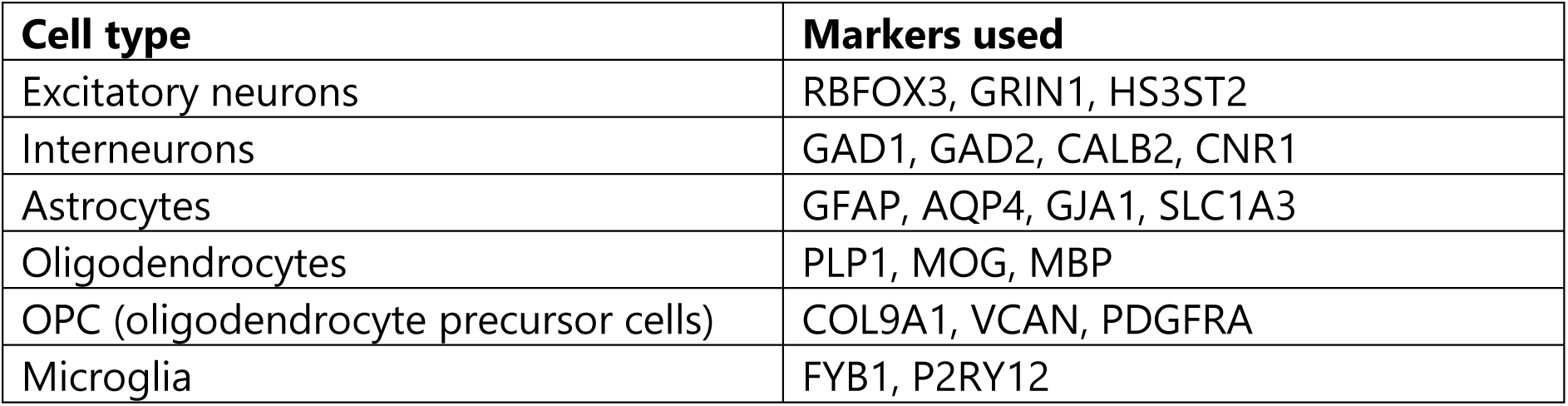

#### Signal of GBA1/GBAP1 per cell type

Barcodes (grouped by sample and cell type) were used to create Cluster objects from the python package trusTEr (version 0.1.1; https://github.com/raquelgarza/truster) and processed with the following functions:

1. tsv_to_bam() – extracts the given barcodes from a sample’s BAM file (outs/possorted_genome_bam.bam output from Cell Ranger count) using the subset-bam software from 10x Genomics (v 1.0). Outputs one BAM file for each cell type per sample, which contains all alignments.
2. filter_UMIs() – filters BAM files to only keep unique combinations of cell barcodes, UMI, and sequences.
3. bam_to_fastq() – uses bamtofastq from 10x Genomics (version 1.2.0) to outputs the filtered BAM files as fastQ files.
4. concatenate_lanes() – concatenates the different lanes (as output from bamtofastq) from one library and generates one FASTQ file per cluster.
5. merge_clusters() – concatenates the resulting FASTQ files (one for each cell type and sample) in defined groups of samples. Here, groups were set to PD or Control depending on the diagnosis of the individual from which the sample was derived. Output is a FASTQ file per cell type per condition.
6. map_clusters() – the resulting FASTQ files were then mapped using STAR (v 2.7.8a). Multimapping reads were allowed to map up to 100 loci (outFilterMultimapNmax 100, winAnchorMultimapNmax 200), the rest of the parameters were used as default.

The resulting BAM files were converted to bigwig files using bamCoverage and normalized by the number of nuclei per group (expression was multiplied by a scale factor of 1e+07 and divided by the number of nuclei in a particular cell type) (deeptools v 2.5.4; RRID:SCR_016366).

For more details, please refer to the scripts process_celltypes_control_PFCTX.py, celltypes_characterization_PFCTX_Ctl.Rmd, and Snakefile_celltypes_control_PFCTX at the github https://github.com/raquelgarza/GBA_snRNAseq_cutnrun_Gustavsson2022.git.

#### CUT&RUN

Post-mortem brain tissue from control individuals with no known history of neurological or neuropsychiatric symptoms was acquired from the Skåne University Hospital Tissue Bank (ethical approvement Ethical Committee in Lund, 06582-2019 & 00080-2019). From the flash-frozen tissue, 50-100 mg were sampled from the dorsolateral prefrontal cortex and stored at −80 °C until use.

CUT&RUN was performed as previously described ^74^, with minor modifications. ConA-coated magnetic beads (Epicypher) were activated by washing twice in bead binding buffer (20 mM HEPES pH 7.5, 10 mM KCl, 1 mM CaCl, 1 mM MnCl_2_) and placed on ice until use. For adult neuronal samples, nuclei were isolated from frozen tissue as described above (see, “Nuclei extraction of cortical post-mortem tissue”). Prior to FACS, nuclei were incubated with Recombinant Alexa Fluor® 488 Anti-NeuN antibody [EPR12763] - Neuronal Marker (ab190195) at a concentration of 1:500 for 30 minutes on ice. The nuclei were run through the FACS at 4 °C at a low flowrate, using a 100 μm nozzle. 300,000 Alexa Fluor – 488 positive nuclei were sorted. The sorted nuclei were pelleted at 1,300 x g for 15 min and resuspended in 1 mL of ice-cold nuclear wash buffer (20 mM HEPES, 150 mM NaCl, 0.5 mM spermidine, 1x cOmplete protease inhibitors, 0.1% BSA). 30 µL (10 µL per antibody treatment) of ConA-coated magnetic beads (Epicypher) were added during gentle vortexing (pipette tips for transferring nuclei were pre-coated with 1% BSA). Binding of nuclei to beads proceeded for 10 min at room temperature with gentle rotation, and then bead-bound nuclei were split into equal volumes (corresponding to IgG control and H3K4me3 treatments). After removal of the wash buffer, nuclei were then resuspended in 100 µL cold nuclear antibody buffer (20 mM HEPES pH 7.5, 0.15 M NaCl, 0.5 mM Spermidine, 1x Roche complete protease inhibitors, 0.02% w/v digitonin, 0.1% BSA, 2 mM EDTA) containing primary antibody (rabbit anti-H3K4me3 Active Motif 39159, RRID:AB_2615077; or goat anti-rabbit IgG, Abcam ab97047, RRID:AB_10681025) at 1:50 dilution and incubated at 4 °C overnight with gentle shaking. Nuclei were washed thoroughly with nuclear digitonin wash buffer (20 mM HEPES pH 7.5, 150 mM NaCl, 0.5 mM Spermidine, 1x Roche cOmplete protease inhibitors, 0.02% digitonin, 0.1% BSA) on the magnetic stand. After the final wash, pA-MNase (a generous gift from Steve Henikoff) was added in nuclear digitonin wash buffer and incubated with the nuclei at 4 °C for 1 h. Nuclei were washed twice, resuspended in 100 µL digitonin buffer, and chilled to 0-2 °C in a metal block sitting in wet ice. Genome cleavage was stimulated by addition of 2 mM CaCl_2_ at 0 °C for 30 min. The reaction was quenched by addition of 100 µL 2x stop buffer (0.35 M NaCl, 20 mM EDTA, 4 mM EGTA, 0.02% digitonin, 50 ng/µL glycogen, 50 ng/µL RNase A, 10 fg/µL yeast spike-in DNA (a generous gift from Steve Henikoff)) and vortexing. After 30 min incubation at 37 °C to release genomic fragments, bead-bound nuclei were placed on the magnet stand and fragments from the supernatant purified by a NucleoSpin clean-up kit (Macherey-Bagel). Illumina sequencing libraries were prepared using the Hyperprep kit (KAPA) with unique dual-indexed adapters (KAPA), pooled and sequenced on a Nextseq500 instrument (Illumina).

#### CUT&RUN analysis

Paired-end reads (2×150 bp) were aligned to the hg38 genome using bowtie2^75^ (v 2.3.4.2; RRID:SCR_016368) (--local –very-sensitive-local –no-mixed –no-discordant – phred33 -I 10 -X 700), converted to bam files with samtools^58^ (v 1.4; RRID:SCR_002105), and indexed with samtools^58^ (v 1.9; RRID:SCR_002105).

Normalized bigwig coverage tracks were made with bamCoverage (deepTools ^76^ v 2.5.4; RRID:SCR_016366), with RPKM normalization. For more details, please refer to the pipeline Snakefile_Neun_cutnrun in the github https://github.com/raquelgarza/GBA_snRNAseq_cutnrun_Gustavsson2022.git.

### TRANSLATION OF NOVEL TRANSCRIPTS

#### Structure predictions

Protein sequences of the different isoforms were aligned pairwise to MANE select with BioPython using a BLOSUM62 scoring matrix with gap open penalty of −3 and gap extend penalty of −0.1. pLDDT scores for residues from AlphaFold2 models were extracted and mapped onto the sequence of MANE select according to the alignment. While the structure of the predictions of newly detected isoforms follows mostly the known GBA1 structure a noteworthy breakdown of the confidence score in regions with deletions is visible. This might indicate a conflict between coevolution information and structural templates from dominant isoforms vs. the learned physico-chemical properties of protein structures, which might be *unfavorable* in those regions.

#### Cell culture

H4 cells (ATCC® HTB-148148™) with homozygous knockout of GBA1 (ENSG00000177628) were generated using indels-based CRISPR/Cas9 technology [gRNA 5’-TCCATTGGTCTTGAGCCAAG-3’ (reverse orientation) targeting exon 7] via Horizon Discovery Ltd. Cells were cultured in DMEM supplemented with 10% foetal bovine serum at 37 °C, 5% CO2. Cells were sub-cultured every 3-4 days at a split ratio of 1:6.

#### Cell transfection

Cells were transfected using Lipofectamine 3000 reagent (Invitrogen L3000008) according to manufacturer’s instructions. *GBA1* or *GBAP1* transcripts subcloned in the pcDNA3.1(+)-C-DYK vector were designed using the GenSmart design tool and acquired from GenScript.

#### Western blot

Protein was extracted from whole cells using MSD lysis buffer (MSD R60TX-3) containing 1x cOmplete Mini Protease Inhibitor Cocktail (Roche 11836153001) and 1x PhosSTOP Phosphatase Inhibitor Cocktail (Roche 4906845001). Protein concentration was determined by Bicinchoninic acid (BCA) assay according to manufacturer’s instructions (Pierce 23225). 10-20 µg of protein diluted in NuPAGE™ LDS Sample Buffer (Invitrogen NP0007) and 200 mM DTT was loaded on NuPAGE™ 4-12% Bis-Tris mini protein gels. Gels were run in NuPAGE™ MES SDS Running Buffer (Invitrogen NP0002) at 150V and transferred to 0.2 µm nitrocellulose membranes in Tris-glycine transfer buffer containing 20% MeOH at 30V for 1.5 hrs. Subsequently, membranes were blocked in Intercept Blocking Buffer (LI-COR 927-60001), incubated with primary antibodies overnight at 4 °C, then IRdye-conjugated secondary antibodies before imaging on the LI-COR Biosciences-Odyssey CLx imaging system. Primary antibodies used include mouse anti-FLAG (Sigma F3165), rabbit anti-GBA1 (C-terminal; Sigma G4171) and rabbit anti-GAPDH (Abcam ab9485).

#### GCase activity assay

Cells cultured on a 96-well plate were washed with PBS (no Ca2+, no Mg2+) and harvested in activity assay buffer containing 50 mM citric acid/potassium phosphate pH 5.0-5.4, 0.25% (v/v) Triton X-100, 1% (w/v) sodium taurocholate, and 1 mM EDTA. After a cycle of freeze/thaw and 30 min incubation on ice, samples were centrifuged at 3,500 rpm for 5 min in 4 °C. Supernatant was collected and incubated in 1% BSA and 2 mM 4-methylumbelliferyl-β-D-galactopyranoside (4-MUG, Sigma M3633) for 90 min at 37 °C. The reaction was stopped by addition of 1 M glycine pH 12.5, and fluorescence (Ex 365 nm; Em 445 nm) was measured using SpectraMax M2 microplate reader (Molecular Devices). Enzyme activity was normalised to untransfected controls.

#### Immunofluorescence

Cells cultured on a 96-well plate were fixed in 4% PFA for 10 min, methanol for 10 min, and permeabilized in 0.3% Triton X-100 for 10 min at room temperature. Cells were then blocked in BlockAce blocking reagent (BioRad BUF029) for 60 min then incubated with primary antibodies at 4 °C overnight. Following washing with PBS with 0.1% Tween-20, cells were incubated with Alexa Fluor secondary antibodies and Hoechst nucleic acid stain. Imaging was performed on the Thunder imager (Leica). Primary antibodies used include mouse anti-FLAG (Sigma F3165), mouse anti-GBA1 (Abcam ab55080) and rabbit anti-Cathepsin D (Abcam ab75852).

#### Mass spectrometric analysis of prefrontal cortex proteomes

A public mass spectrometry dataset was retrieved from ProteomeXchange (PXD026370). This data set consists of human brain tissue was collected post-mortem from patients diagnosed with multiple system atrophy (*n* = 45) and from controls (*n* = 30) to perform a comparative quantitative proteome profiling of tissue from the prefrontal cortex (Broadman area 9)^46^.

The data analysis was performed using MetaMorpheus^77^ (v 0.0.320; https://github.com/smith-chem-wisc/MetaMorpheus). The search was conducted for two 2 GBAP1 isoforms (PB.845.1693 and PB.845.525), and a list of 267 frequent protein contaminants found within mass spectrometry data as provided by MetaMorpheus. An FDR (false discovery rate) of 1% was applied for presentation of PSMs (peptide spectrum matches), peptides, and proteins following review of decoy target sequences.

The following search settings were used: protease = trypsin; maximum missed cleavages = 2; minimum peptide length = 7; maximum peptide length = unspecified; initiator methionine behavior = Variable; fixed modifications = Carbamidomethyl on C, Carbamidomethyl on U; variable modifications = Oxidation on M; max mods per peptide = 2; max modification isoforms = 1024; precursor mass tolerance = ±5.0000 PPM; product mass tolerance = ±20.0000 PPM; report PSM ambiguity = True.

### ANNOTATION OF PARENT GENES AND PROTEIN-CODING GENES

To explore inaccuracies in annotation of parent genes and protein-coding genes we applied three independent approaches:

#### Long-read RNA-sequencing

To identify full-length transcripts with at least one novel splice junction we used the same long-read RNA-seq samples available from ENCODE^53^ as previously described. Transcripts with novel splice junction resulting in novel ORF were those transcripts that had a predicted ORF that was not present in GENCODE v38 annotation.

#### Novel expressed regions

Novel unannotated expression^37^ was downloaded from Visualisation of Expressed Regions (vizER; https://rytenlab.com/browser/app/vizER). The data originates from RNA-seq data in base-level coverage format for 7,595 samples originating from 41 different GTEx tissues. Cell lines, sex-specific tissues, and tissues with 10 samples or below were removed. Samples with large chromosomal deletions and duplications or large copy number variation previously associated with disease were filtered out (smafrze = “USE ME”). Coverage for all remaining samples was normalized to a target library size of 40 million 100-bp reads using the area under coverage value provided by recount2^52^. For each tissue, base-level coverage was averaged across all samples to calculate the mean base-level coverage. GTEx junction reads, defined as reads with a non-contiguous gapped alignment to the genome, were downloaded using the recount2 resource and filtered to include only junction reads detected in at least 5% of samples for a given tissue and those that had available donor and acceptor splice sequences.

#### Splice junctions

To identify novel junctions with potential evidence of incomplete annotation, we used data provided by IntroVerse^78^.

IntroVerse is a relational database that comprises exon-exon split-read data on the splicing of human introns (Ensembl v105) across 17,510 human control RNA samples and 54 tissues originally made available by GTEx and processed by the recount3 project^33^. RNA-seq reads provided by the GTEx v8 project were sequenced using the Illumina TruSeq library construction protocol (non-stranded 76bp-long reads, polyA+ selection). Samples from GTEx v8 were processed by recount3 through Monorail (STAR^56^) to detect and summarise splice junctions and Megadepth^79^ to analyse the bam files produced by STAR). Additional quality-control criteria applied by IntroVerse included: (i) exclusively analysing samples passing the GTEx v8 minimum standards (smafrze != “EXCLUDE”); (ii) discarding any split-reads overlapping any of the sequences included in the ENCODE Blacklist^80^; (iii) or split reads that presented an implied intron length shorter than 25 base pairs.

Second, we extracted all novel donor and acceptor junctions that had evidence of use in >=5% of the samples of each tissue and grouped them by gene. We then classify those genes either as “parent” or “protein-coding.” Finally, we calculated the proportion that each category of genes presented within each tissue. Focusing on the *parent genes* category, this can be described as it follows:

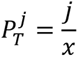

Let *j* denote the total number of parent genes containing at least one novel junction shared by >=5% of the samples of the current tissue. Let *x* denote the total number of *parent* genes available for study. Let *T* denote the current tissue.

We mirrored the formula above to calculate the proportion of protein-coding genes per tissue.

## FIGURE GENERATION

The code for all figures in this manuscript can be accessed through: https://github.com/egustavsson/GBA_GBAP1_manuscript.git

## ACKNOWLEDGMENTS

This research was funded in whole or in part by Aligning Science Across Parkinson’s [Grant numbers: ASAP-000478, ASAP-000509, and ASAP-000520] through the Michael J. Fox Foundation for Parkinson’s Research (MJFF). For the purpose of open access, the author has applied a CC BY public copyright licence to all Author Accepted Manuscripts arising from this submission.

E.K.G. was also supported by the Postdoctoral Fellowship Program in Alzheimer’s Disease Research from the BrightFocus Foundation (Award Number: A2021009F). C.H.D. was supported by a Swedish Society for Medical Research Starting Grant (SSMF S19-0100). M.R. was supported through the award of a Tenure Track Clinician Scientist Fellowship (MR/N008324/1). This work was funded by a postdoctoral fellowship awarded to S.S. and Y.G. under the “Sustaining Innovation Postdoctoral Training Programme” at Astex Pharmaceuticals. C.A. is supported by a fellowship from the Alzheimer’s Society (AS-JF-18-008). S.W. is supported by an Alzheimer’s Research UK Senior Research Fellowship (ARUK-SRF2016B-2). S.W., C.A. and J.H. are supported by the NIHR UCL Hospitals Biomedical Research Centre.

## COMPETING INTERESTS

S.S., Y.G., J.E., H.S. and C.F.B. are employed by Astex Pharmaceuticals. The other authors declare no competing interests.

## EXTENDED DATA

**Extended Data Fig. 1:**
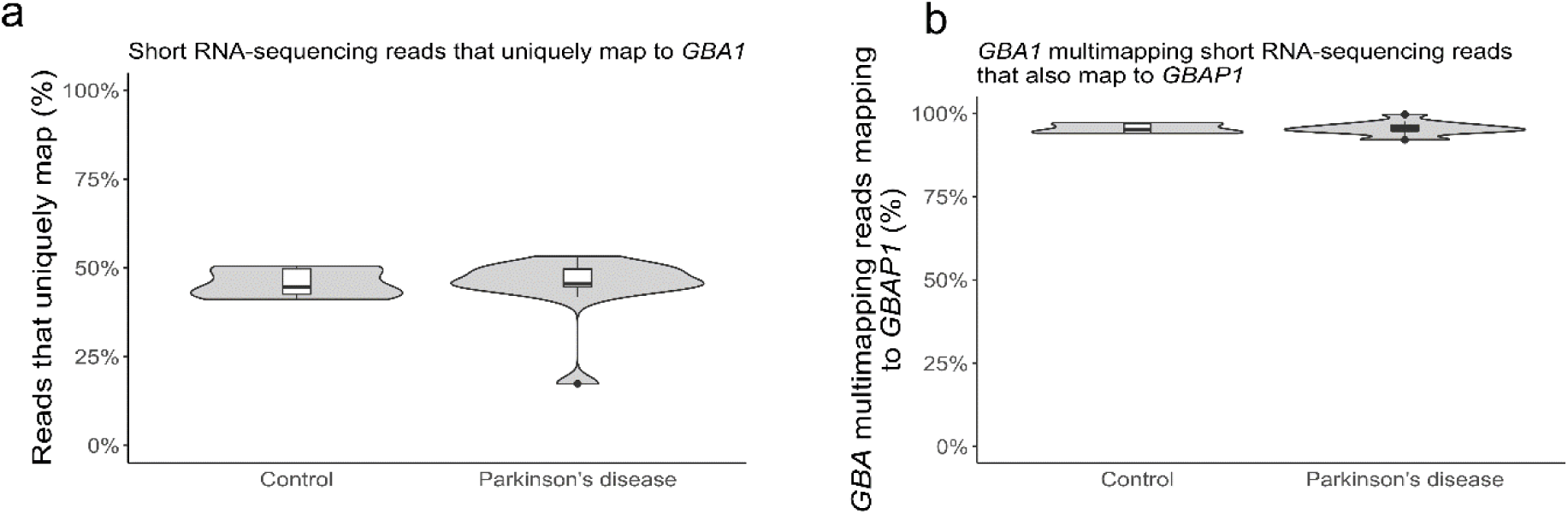
Most short RNA seq reads mapping to *GBA1* multimap to *GBAP1*. **a,** Violin plots showing multimapping of *GBA1* from short-read RNA-seq data (100bp paired end reads, mean reads per sample of 182.9 ± 14.9M) from human post-mortem anterior cingulate cortex samples generated from control (*n* = 5) and PD-affected individuals (*n* = 7)^34^. **b,** Violin plots showing the percentage of *GBA1* short RNA-sequencing multimapping reads that that also map to *GBAP1*.

**Extended Data Fig. 2:**
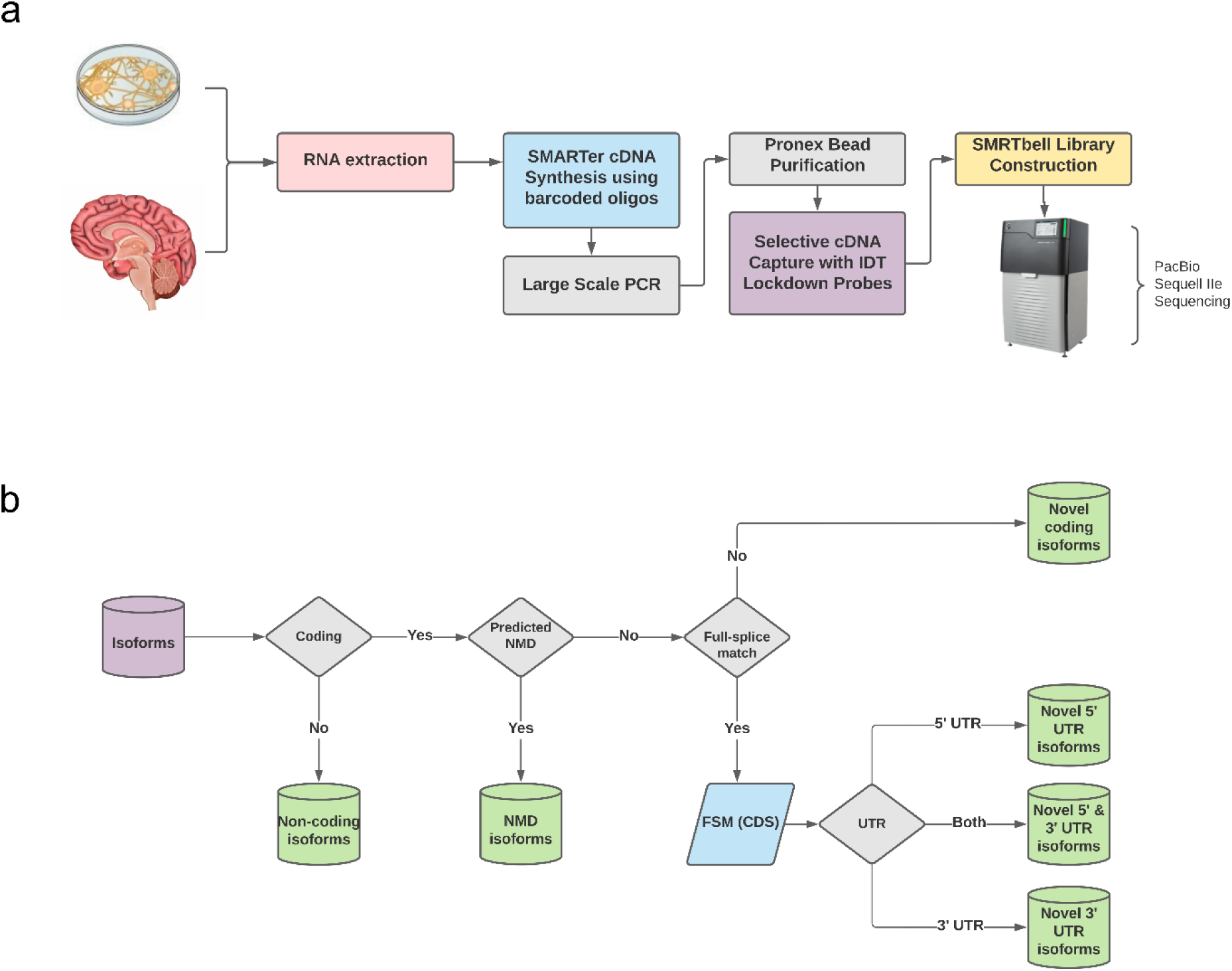
Approach for targeted long-read RNA-sequencing. **a,** Schematic illustration showing the approach taken for targeted long-read RNA-sequencing of *GBA1* and *GBAP1* in human brain tissues and iPSC derived neurons, microglia, and astrocytes. **b,** Flowchart showing the categorization of transcripts generated though long-read RNA-sequencing.

**Extended Data Fig. 3:**
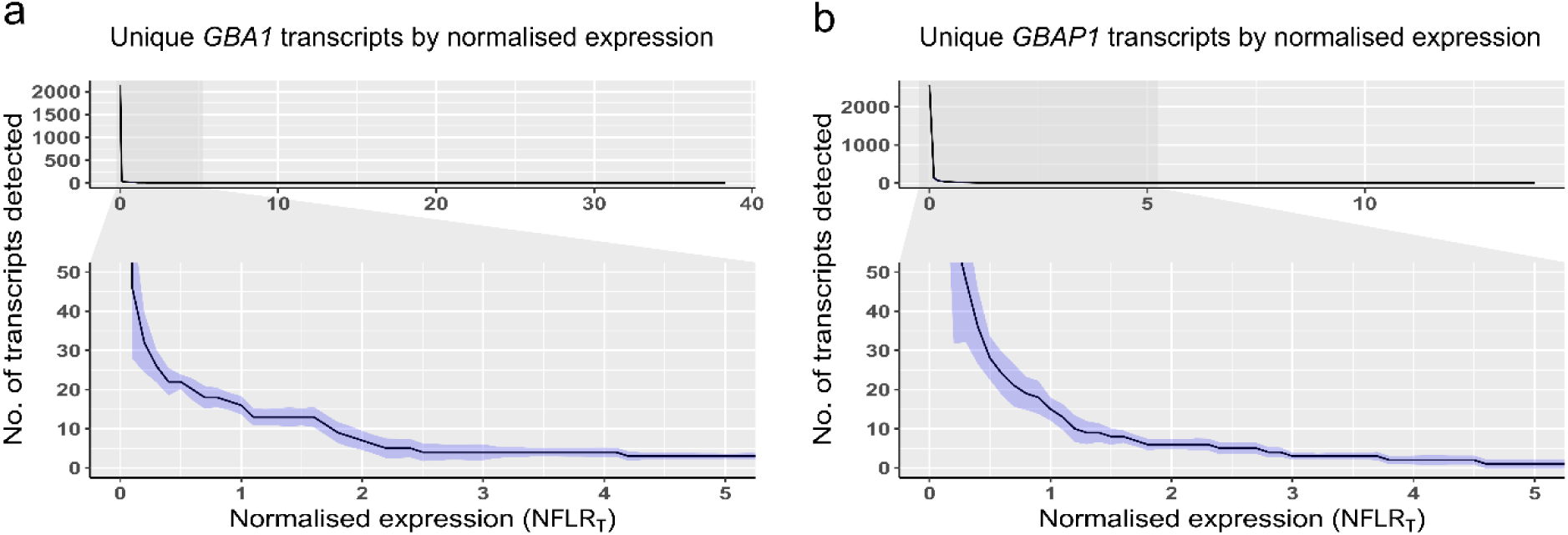
Total number of unique transcripts of *GBA1* and *GBAP1* by normalized expression. **a,** Depreciation curve showing the number of unique *GBA1* transcripts on the Y-axis increased by increasing the normalized full-length read count of transcript (NFLR_T_) on the X-axis. NFLR_T_ is the total number of reads per transcript normalized by the total number of reads of the loci. **b,** Depreciation curve showing the number of unique *GBAP1* transcripts on the Y-axis increased by increasing the NFLR_T_ on the X-axis.

**Extended Data Fig. 4:**
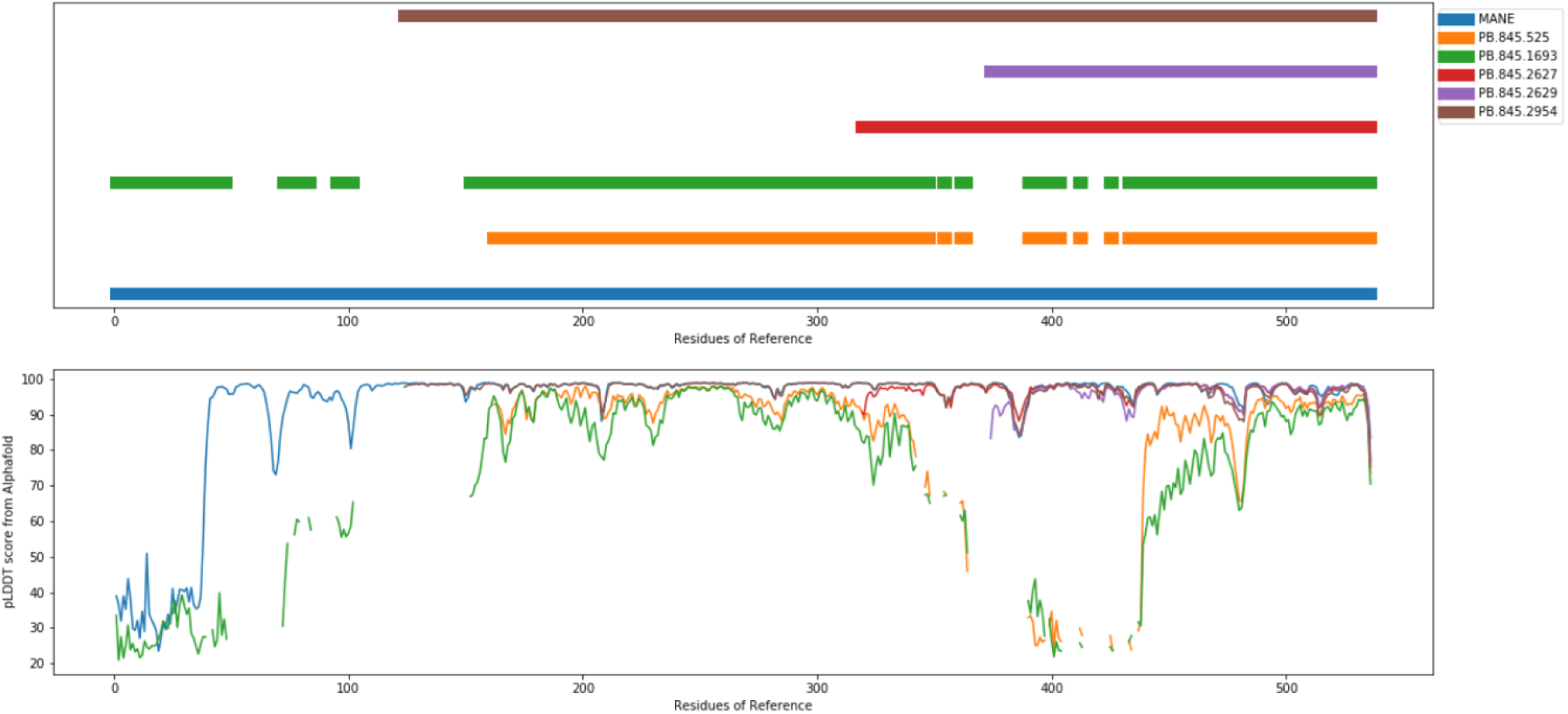
Alignment of novel GBA1 and GBAP1 protein sequences. Protein sequences of novel GBA1 and GBAP1 isoforms pairwise aligned to GBA1 MANE select.

**Extended Data Fig. 5:**
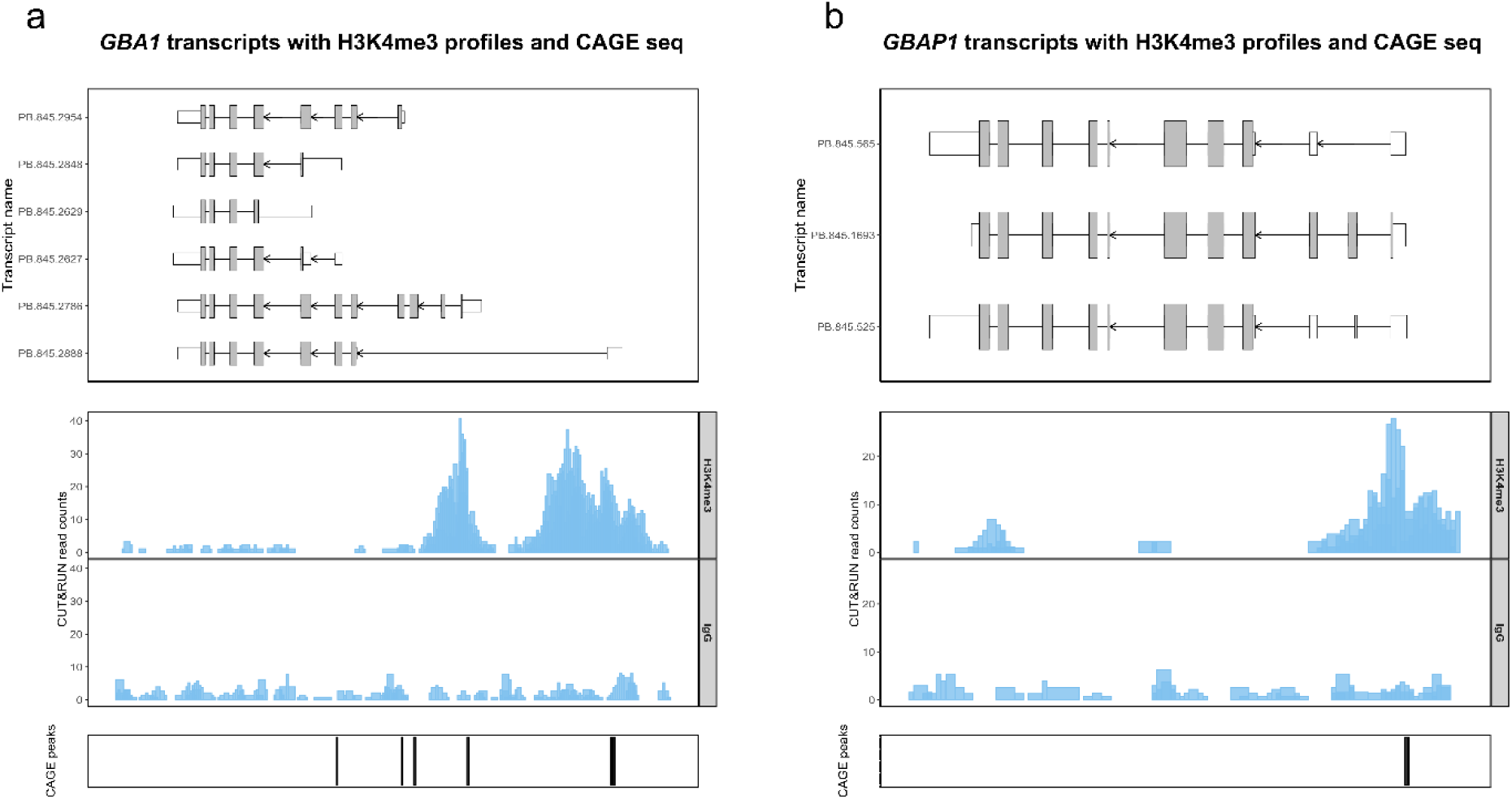
Transcriptionally active euchromatin at the 5’ TSS of GBAP1 ORF transcripts. **a,** Novel protein-coding transcripts of *GBA1* CUT&RUN profiling of H3K4me3 marks in neurons (based on NeuN+) and CAGE sequencing data from FANTOM5. **b,** Novel protein-coding transcripts of *GBAP1* CUT&RUN profiling of H3K4me3 marks in neurons (based on NeuN+) and CAGE sequencing data from FANTOM5.

## Notes

### Summary of Updates

This version includes an updated title, introduction and discussion. It also include additional untargeted long-read RNA-seq data for the analysis of novel splicing of parent genes and alternative splicing of pseudogenes.

