## supplementary figures for "The annotation and function of the Parkinson’s and Gaucher disease-linked gene *GBA1* has been concealed by its protein-coding pseudogene *GBAP1*"

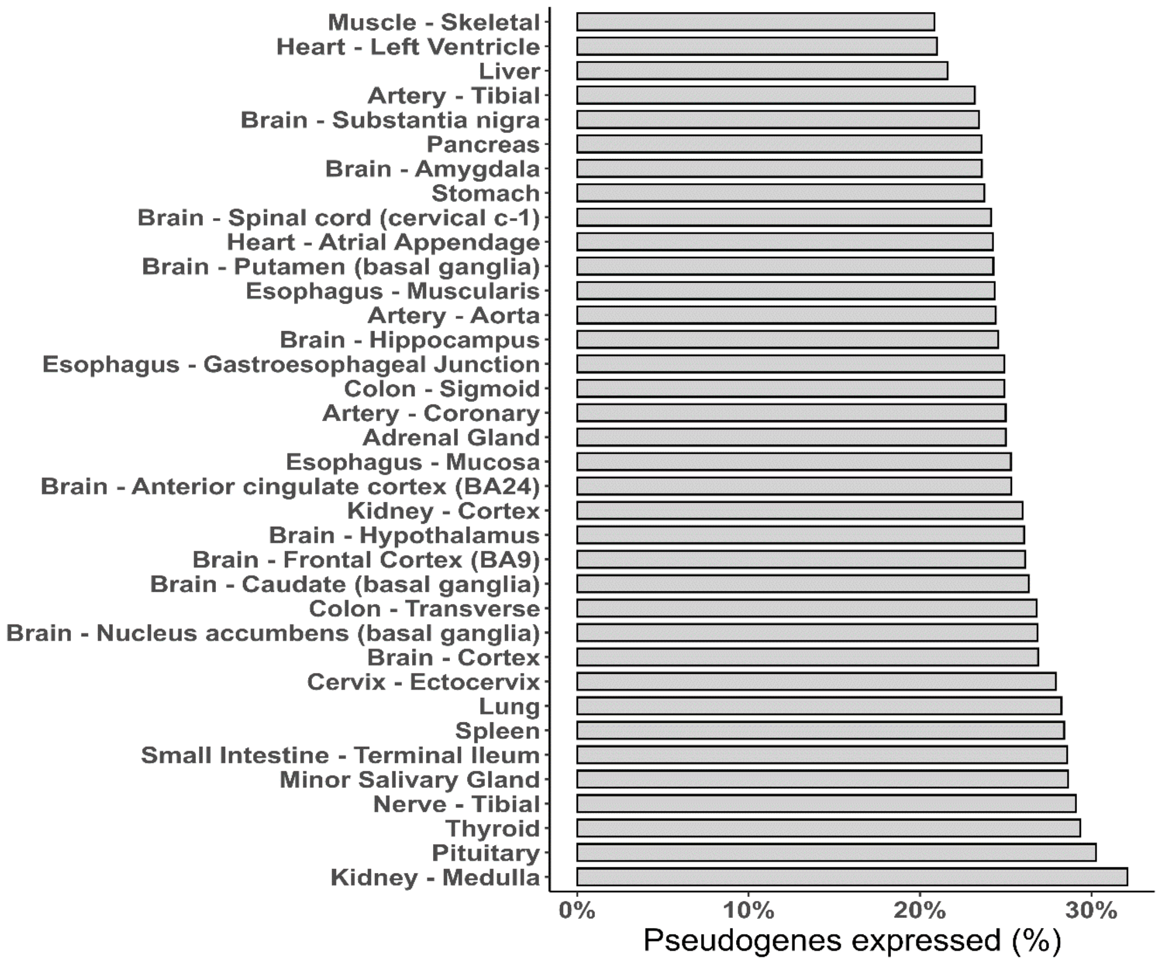


**Supplementary Fig. 1: Pseudogenes are frequently expressed across human tissues.** Histogram showing the percentage of human pseudogenes expressed per tissues as assessed using uniquely mapping reads (generated by the Genotype-Tissue Expression Consortium, GTEx v8). A gene with a median TPM > 0 was considered to be expressed.

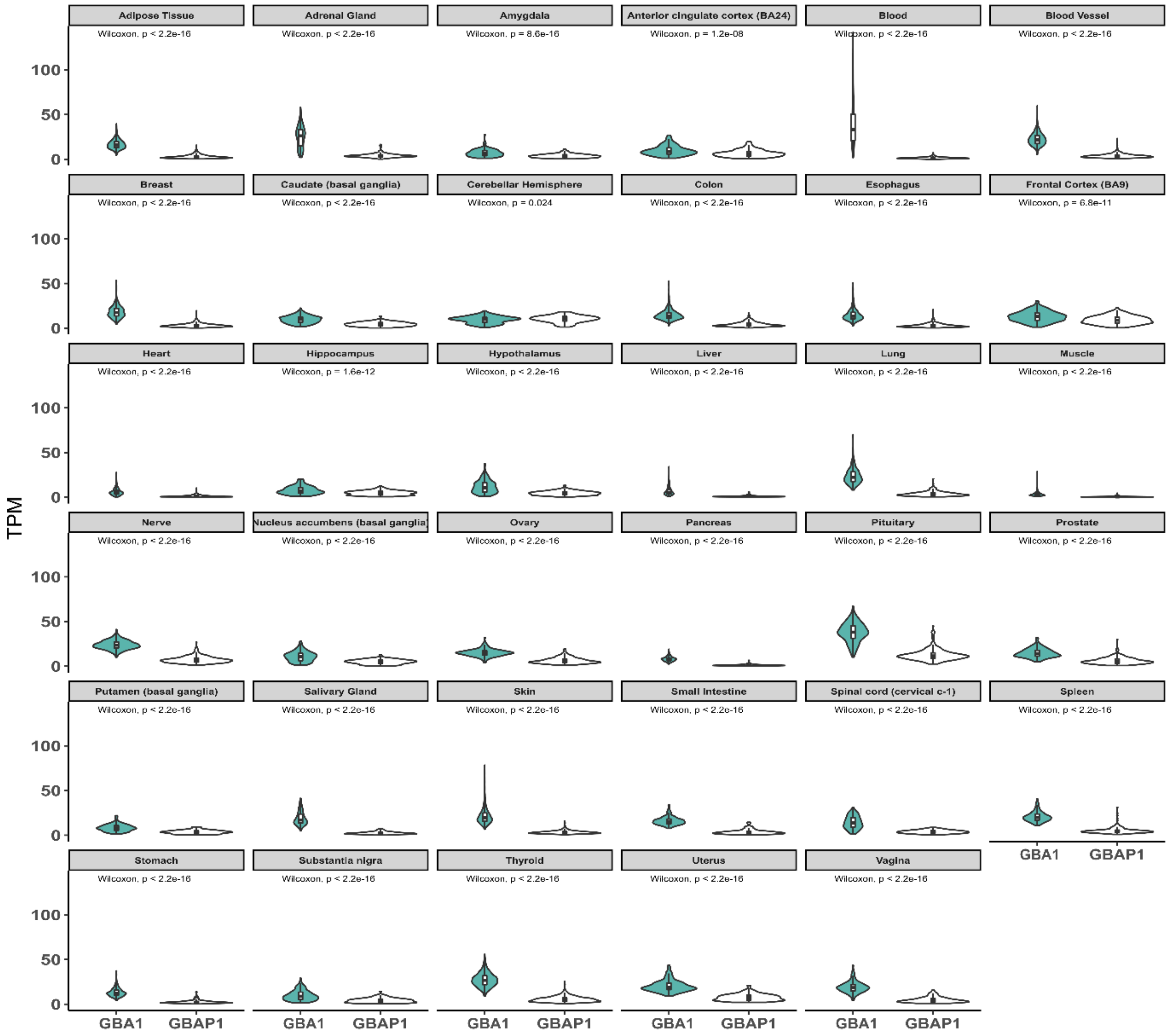


**Supplementary Fig. 2: *GBA1* and *GBAP1* are widely expressed across human tissues.** Violin plots showing the transcripts per million (TPM) expression of *GBA1* and *GBAP1* across human tissues generated by the Genotype-Tissue Expression Consortium, GTEx v8**).**

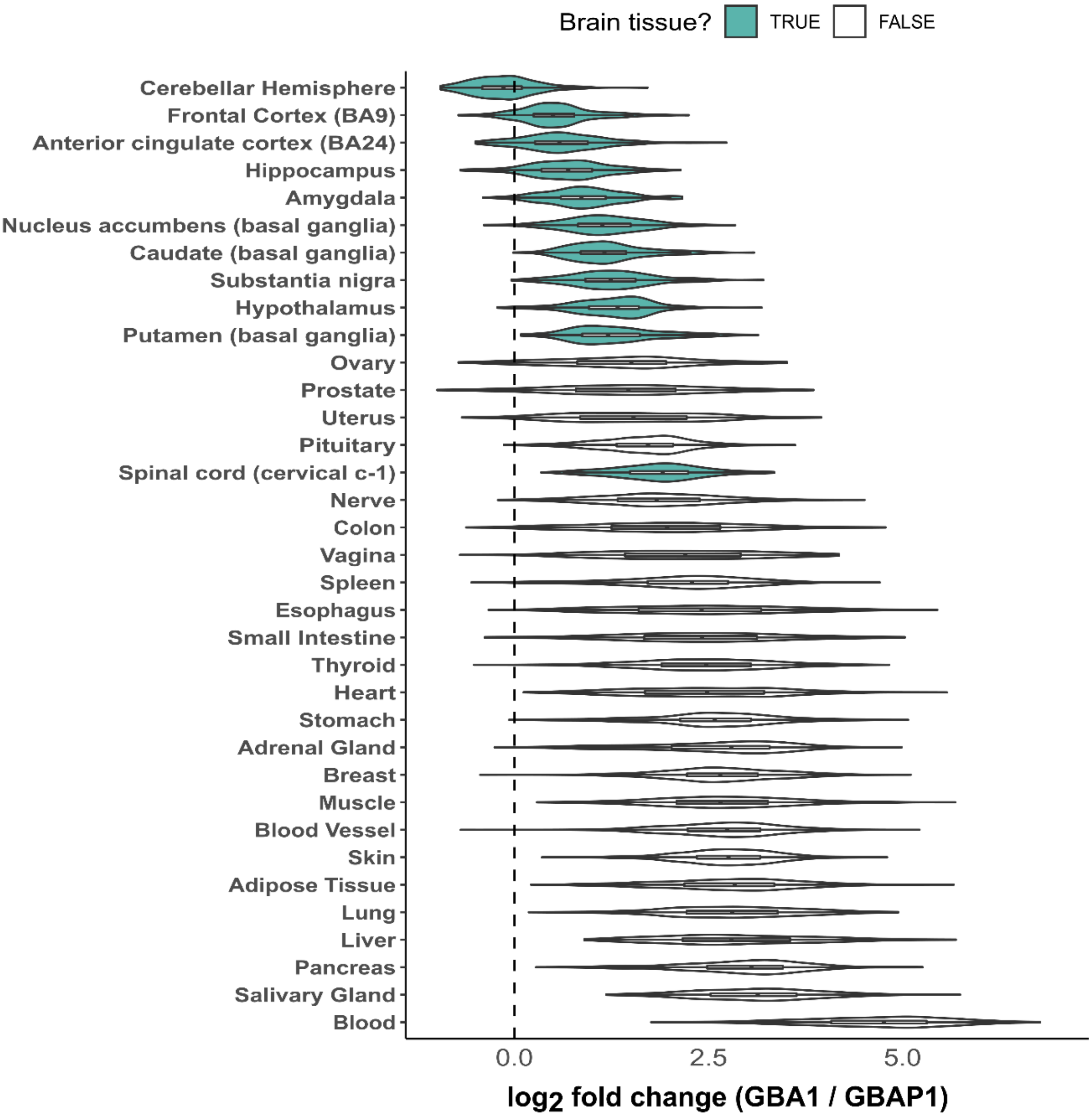


**Supplementary Fig. 3: Expression of *GBAP1* is similar to *GBA1* in human brain.** Violin plots showing $log_2$ fold change of *GBA1* (numerator) by *GBAP1* (denominator) across human tissues generated by the Genotype-Tissue Expression Consortium, GTEx v8**).**

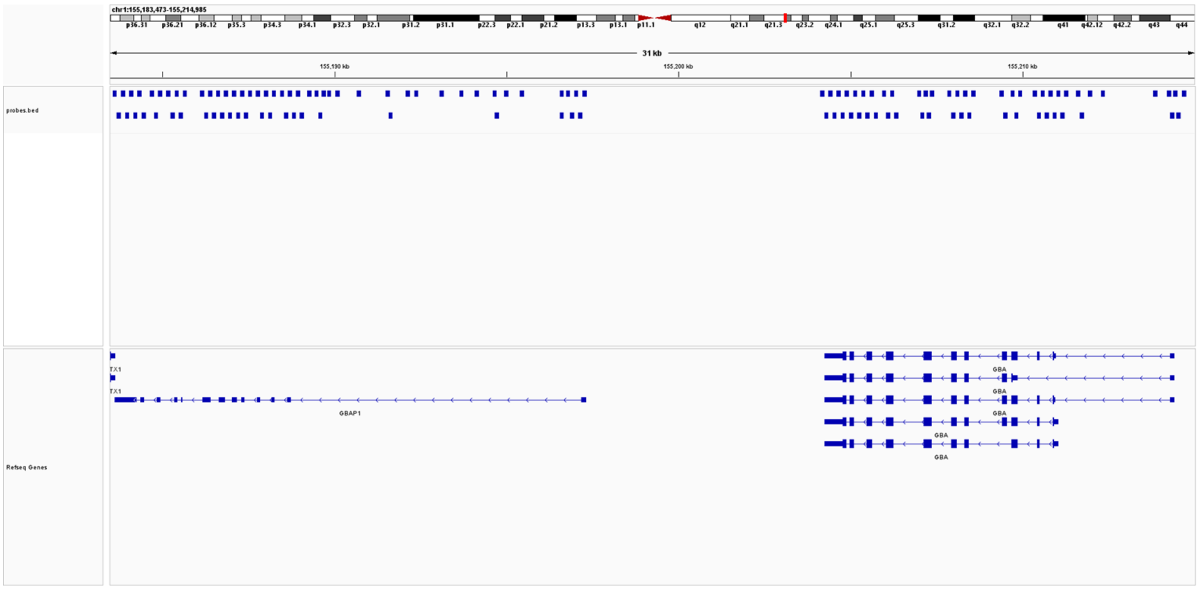


**Supplementary Fig. 4: *GBA1* and *GBAP1* hybridization probe design.** 120-mer IDT lockdown hybridization probe design used for the enrichment of *GBA1* and *GBAP1* cDNA.

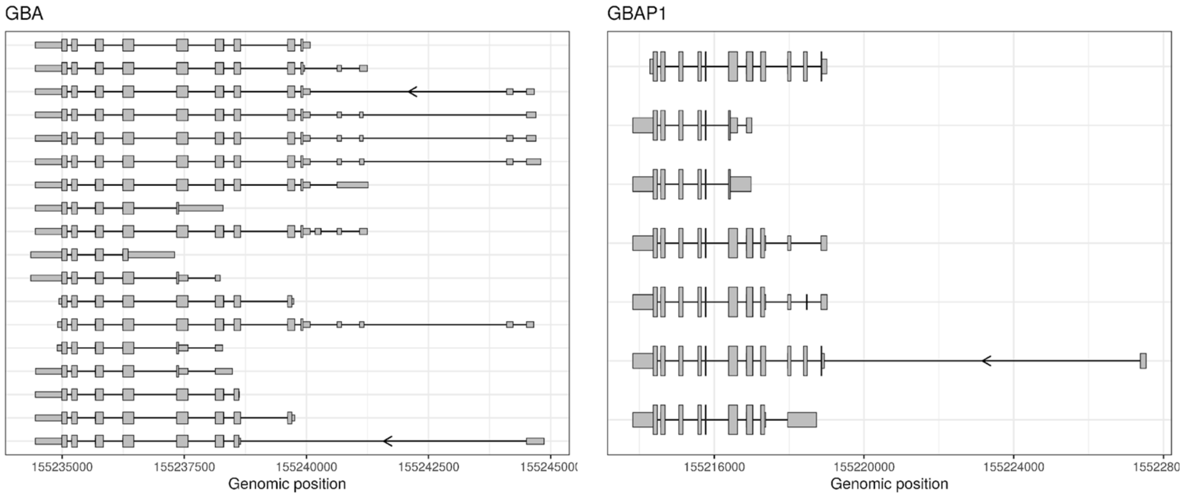


**Supplementary Fig. 5: *GBA1* and *GBAP1* transcripts with novel open reading frames. a,** 18 novel *GBA1* transcripts with novel open reading frames (ORF) identified through targeted long-read RNA sequencing of 12 human brain regions. **b,** 7 *GBAP1* transcripts with ORFs predicted to be coding identified through targeted long-read RNA sequencing of 12 human brain regions.

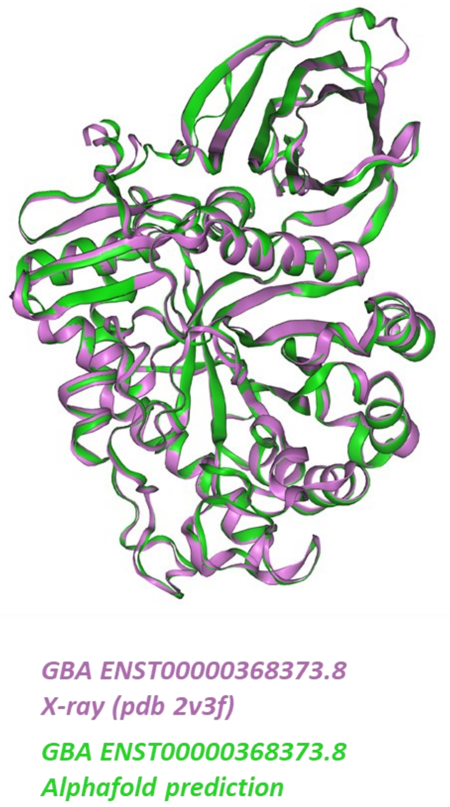


**Supplementary Fig. 6:** Experimental X-ray structure of MANE select (PDB ID 2v3f) (violet) superimposed on AlphaFold2 prediction of the same sequence (green).

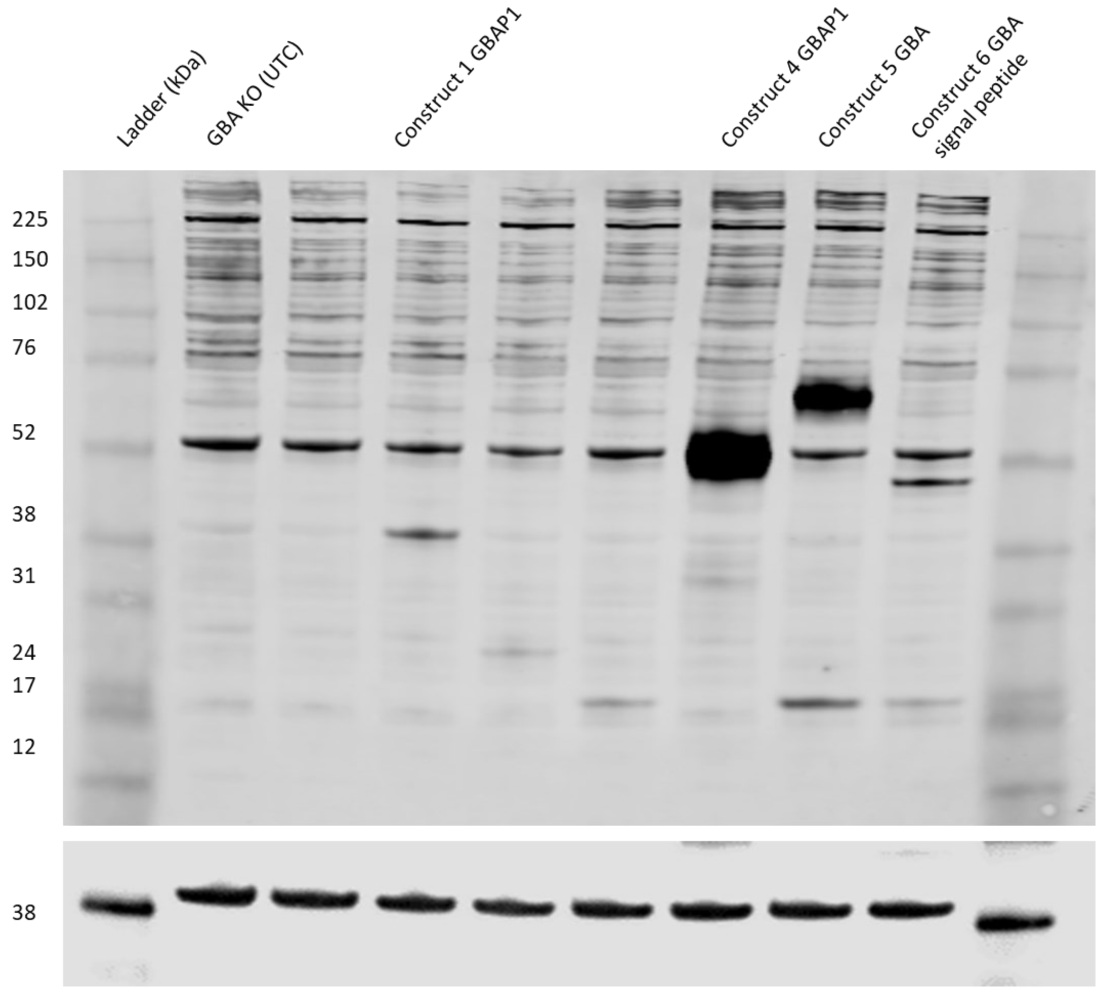


**Supplementary Fig. 7:** Immunoblot of H4 GBA1(-/-/-) knockout cells transiently transfected with GBA1 and GBAP1 constructs containing a c-terminus FLAG-tag. GBA1 and GBAP1 expression was detected using GBA1 anti-rabbit G4171 from Sigma-Aldrich (1:1000), synthetic peptide corresponding to amino acids 517-536 (c-terminus) of human glucocerebrosidase conjugated to KLH. GAPDH was used as a loading control. The predicted protein sizes are: PB.845.525 (GBAP1; 321 aa; 35 kDa), PB.845.2627 (GBA1 affecting GH30 and SP; 219 aa; 24 kDa), PB.845.2629 (GBA1 affecting GH30 and SP; 164 aa; 18 kDa), PB.845.1693 (GBAP1; 399 aa; 44 kDa), ENST00000368373 (GBA1 MANE select; 537 aa; 62 kDa) and PB.845.2954 (GBA1 affecting GH30 and SP; 414 aa; 46 kDa).

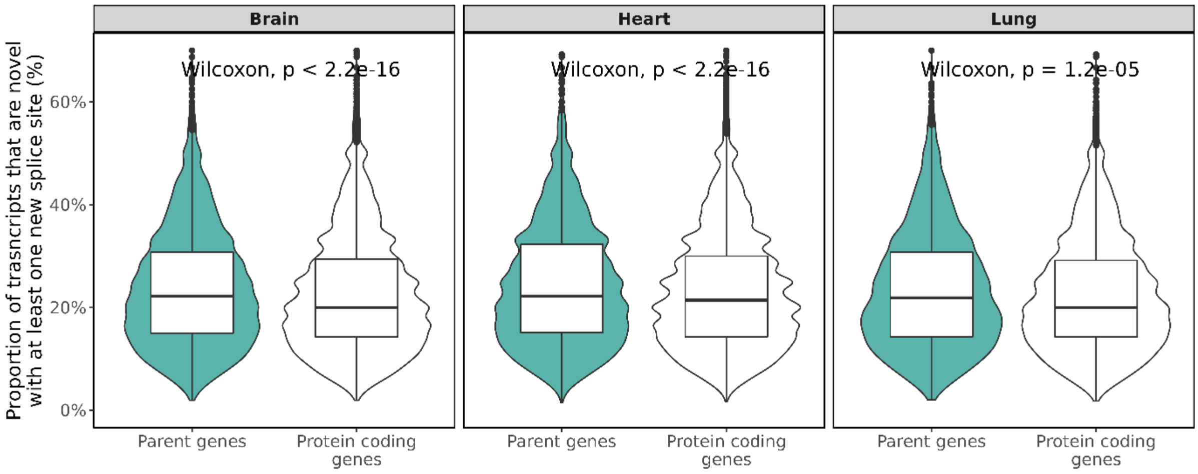


**Supplementary Fig. 8:** Proportion of transcripts per parent gene and per protein coding gene without a pseudogene with a novel splice site from long-read RNA-sequencing data in Brain (*n* = 9), Heart (*n* = 16) and lung (*n* = 6).
